# Systematic Quantitative Proteomics Defines Age, Sex, and Region-Dependent Remodeling of Lung Extracellular Matrix

**DOI:** 10.64898/2026.06.19.733262

**Authors:** Anna G. Towler, Fei Wang, Yingtao Bi, Liam J. Bandura, Yanlong Zhu, Junjie Zhu, Andrew J. Perciaccante, Timothy J. Aballo, Qin C. Ji, Liang Jin, Wanye Buck, Lucy Phillips, Kuniko Kadoya, Lynn M. Schnapp, Yupeng He, Yu Tian, Ying Ge

**Affiliations:** Department of Chemistry, University of Wisconsin-Madison, Madison, WI 53706; Quantitative Translational & ADME Science, AbbVie Bioresearch Center, Worcester, MA 01605; Department of Cell and Regenerative Biology, School of Medicine and Public Health, University of Wisconsin-Madison, Madison, WI 53705; Human Proteomics Program, School of Medicine and Public Health, University of Wisconsin-Madison, Madison, WI 53705; Quantitative Translational & ADME Science, AbbVie Inc., North Chicago, IL 60064; Molecular and Cellular Pharmacology Training Program, University of Wisconsin-Madison, Madison, WI 53705; Discovery Immunology, Pharmacology and Pathology, AbbVie, Inc., North Chicago, IL 60064; Allergan Aesthetics, an AbbVie company, 2525 Dupont Drive, Irvine, CA 92612; Department of Medicine, School of Medicine and Public Health, University of Wisconsin-Madison, Madison, WI 53705

**Author notes:** These two authors contribute equally to this study (Anna G. Towler, Fei Wang). To whom correspondence should be addressed: Dr. Ying Ge, 1111 Highland Ave., WIMR II 8551, Madison, WI 53705. Dr. Yu Tian, 100 Research Dr. Worcester, MA, 01605. Dr. Yupeng He, 1 North Waukegan Road, North Chicago, IL 60064. Dr. Lynn Schnapp, 1685 Highland Ave, Suite 5000, Madison, WI 53705.

**Keywords:** extracellular matrix, matrisome, mass spectrometry, proteomics, aging, sex, anatomical region, collagen crosslink

## Abstract

The lung extracellular matrix (ECM) governs tissue architecture, mechanics, and function, yet how it remodels with age across sex and anatomical regions remains poorly understood. Here, we performed a systematic multi-factor proteomic analysis of rat lungs to define age-, sex-, and region-dependent remodeling across the tissue landscape. Age emerged as the dominant source of variation, with a conserved aging signature modified by region- and sex-specific effects. Young lungs showed coordinated ECM assembly, balanced proteolysis, and active biosynthetic programs consistent with structural adaptability and mechanical resilience. In contrast, aged lungs exhibited accumulation of mature collagen crosslinks and a more stabilized matrix architecture, indicating progressive matrix maturation and reduced structural plasticity. These changes were accompanied by proteomic signatures of metabolic stress and immune activation, suggesting coordinated remodeling across ECM, metabolic, and immune pathways during lung aging. Aging effects varied across anatomical regions and were more pronounced in females, highlighting context-dependent trajectories within the broader aging program. Age also partially reshaped spatial proteomic heterogeneity across lung compartments. Together, these findings identify matrix stabilization as a central feature of lung aging that links structural remodeling to metabolic-inflammatory imbalance and increased pulmonary vulnerability.

## Introduction

The extracellular matrix (ECM) is a dynamic and highly organized network of glycoproteins, proteoglycans, and collagens that governs tissue architecture and mechanical function.^1–3^ By providing both structural support and biochemical/biomechanical signaling cues, the ECM regulates essential cellular processes including migration, proliferation, and differentiation to maintain tissue homeostasis.^1,3,4^ Proper ECM function depends on tightly controlled composition, organization, and remodeling; disruption of these processes alters tissue mechanics, often resulting in increased stiffness and functional decline.^3,5^

In the lung, ECM composition and architecture are uniquely adapted to support repetitive cycles of mechanical stretch during respiration.^6,7^ Fibrillar proteins such as collagens and elastin provide tensile strength and elastic recoil, while ECM crosslinking further modulates tissue stiffness by stabilizing protein–protein interactions. This precise balance between elasticity and structural integrity is essential for maintaining lung compliance and efficient ventilation. Consequently, disruption of ECM composition or organization alters lung physiological function and contributes to pulmonary dysfunction, particularly in the context of aging and pathological remodeling.^8,9^

Aging is a major driver of lung remodeling and is associated with progressive alterations in tissue structure and homeostasis. These changes are accompanied by shifts in ECM composition, tissue mechanics, and molecular regulation, reflecting coordinated remodeling across structural and biochemical systems.^10,11^ While prior transcriptomic and proteomic studies in human and model organisms have characterized age-associated changes in cellular programs and ECM regulation, how aging reshapes the lung proteome at the tissue level remains incompletely understood.^12,13^

In addition to age, biological sex and anatomical region represent important sources of variability in lung structure and molecular composition.^14,15^ Sex-dependent differences influence susceptibility and progression across lung conditions, suggesting underlying divergence in tissue organization and regulatory pathways.^16,17^ Furthermore, the lung exhibits pronounced spatial heterogeneity, with distinct anatomical regions experiencing different mechanical environments and microstructural demands. Recent studies have highlighted region-specific variation in gene and protein expression across the lung, underscoring the importance of spatial context in defining tissue biology.^18,19^

Despite these advances, the combined influence of age, sex, and anatomical region on the baseline lung proteome has not been systematically defined. Here, we performed a systematic proteomic analysis of the rat lung to investigate how these key physiological variables collectively shape the pulmonary proteome, with a particular focus on extracellular matrix components. The rat model enables reproducible regional dissection and provides sufficient tissue for spatially resolved proteomic analysis. We employed a high-throughput, single-step Azo-enabled ECM protein extraction method coupled with bottom-up mass spectrometry to enable comprehensive profiling across this multi-factor design. Our analysis confirms aging as the dominant driver of proteomic remodeling, characterized by a shift from structural maintenance toward immune- and stress-associated programs, while sex and anatomical region modulate the magnitude and spatial organization of these changes.

## Results

### Comprehensive proteomic profiling enables multidimensional analysis of age, sex, and region-dependent remodeling

To define how age, sex, and anatomical region shape the lung proteome, we performed a large-scale mass spectrometry analysis of lung tissue using our previously validated single-step ECM extraction workflow.^20,21^ This approach provides coverage comparable to the dual-step method while substantially improving throughput, enabling systematic profiling of age-, sex-, and region-dependent proteomic remodeling with particular emphasis on ECM components (**Fig. 1**). We analyzed 12 young (7 weeks) and 12 aged (7 - 11 months) rats, including 6 females and 6 males in each age group (**Table S1**). The right lungs were used for H&E histological evaluation as quality control (**Fig. S1a**). Most of the aged rats showed mild macrophage infiltration in the subpleural region of lung lobes, but no tumors were observed. Because the left rat lung contains only a single lobe, we sampled distinct regions (bottom, central, top and peripheral) within the left lung to evaluate regional variation while reducing potential confounding from lobar developmental differences (**Fig. S1b**). Rats are supine animals, so ‘top’ is used to describe the rostral region, and ‘bottom’ refers to the caudal region. In total, 96 freshly frozen lung samples were collected across 24 rats (6 young females, 6 young males, 6 aged females, and 6 aged males) with four separate regions from each set of rat lungs. To facilitate comparisons across sample groups, a standardized nomenclature was adopted; for example, tissue from the bottom (B) region of an aged (A) female (F) rat is denoted as AFB, with similar abbreviations applied to all 16 sample groups (**Table S1**).

**Fig. 1:**
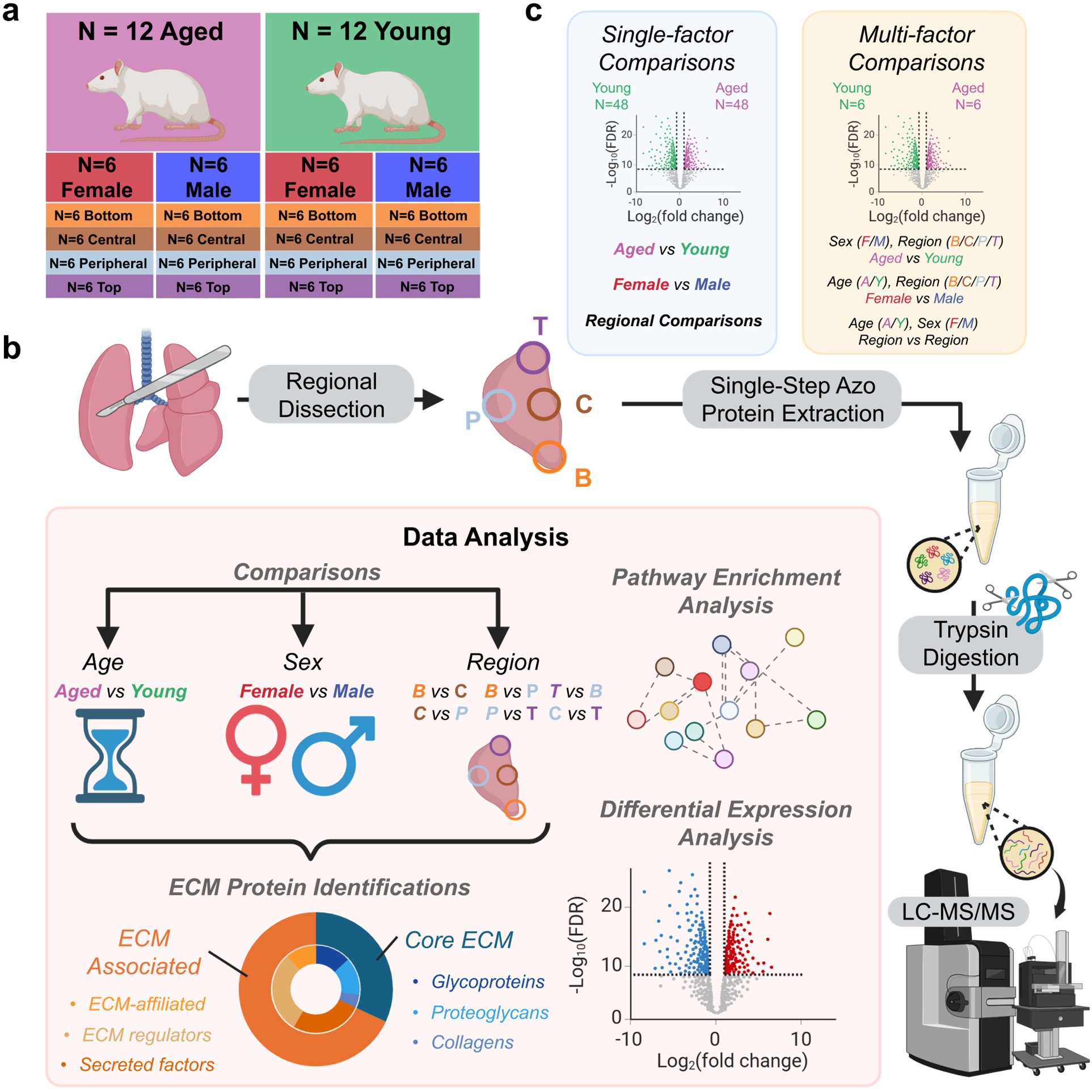
Study design and quantitative proteomics workflow for analysis of rat lung aging, sex, and regional heterogeneity. **a** Overview of the rat lung cohort for proteomics analysis. Sprague–Dawley (SD) rats were grouped by age and sex (12 aged (A): 6 female (F), 6 male (M); 12 young (Y): 6 female, 6 male). From each rat, the left lung was dissected into four anatomical regions—bottom (B), central (C), peripheral (P), and top (T)—yielding 48 young and 48 aged samples. **b** A quantitative proteomics workflow was implemented to examine age, sex, and region-dependent remodeling in rat lungs. **c** Single-factor and multi-factor comparisons were performed to study the effect of age, sex, and/or region on matrisome protein expression. Single-factor comparisons included all young with aged tissues, all female versus all male tissues, and pairwise regions to one another. Multi-factor comparisons similarly compared young vs aged samples, for example, but separated groups based on sex and region while analyzing the effect of age on the rat lung proteome.

The single-step procedure demonstrated high technical reproducibility across biological replicates, with coefficient of variation values (CV) consistently less than 5% (**Fig. S2a**). In total, we identified 3,859 rat protein groups (unfiltered) from 96 individual samples categorized into 16 experimental groups, searching against the Swiss-Prot reviewed UniProtKB reference rat proteome with 8,202 FASTA entries (**Table S2, S3**). Across biological replicates, each sample type yielded between 2,984 and 3,600 protein identifications, except for one AMT sample showing a lower count (2,269 proteins; **Fig. S2b**). On average, 31,387 ± 4,514 (SD) peptides were detected across all samples (**Fig. S2c**).

Gene symbols of identified proteins were searched against the Matrisome AnalyzeR database to annotate the total number of rat matrisome proteins.^22^ On average, the study detected 162 ± 6.87 matrisome proteins per sample, comprising 52.4 ± 2.00 core matrisome (**Fig. S2d, e**) and 110 ± 5.40 (SD) matrisome-associated proteins (**Fig. S2f, g**). When combining all samples before filtering, a total of 185 unique matrisome proteins were identified, including 60 core and 125 matrisome-associated proteins (**Fig. S2h**). Core matrisome proteins included collagens, elastin, and fibronectin, and matrisome-associated proteins included surfactant proteins, matrix metalloproteinases (MMPs), and procollagen-lysine, 2-oxoglutarate 5-dioxgenase (PLOD) hydroxylation enzymes (**Fig. S2h**).

After removing low-recurrence protein groups (see Methods), a filtered dataset of 3,600 protein groups with 8.5% missing values was retained for quantification. Missing data were imputed using a two-step approach — random forest imputation followed by conditional quantile regression imputation of left-censored (QRILC) values, to generate a complete abundance matrix with comprehensive matrisome coverage, enabling robust quantitative analyses of age-, sex-, and region-dependent remodeling (**Table S4**).

Given the hierarchical design incorporating three biological factors: age (aged vs young), sex (female vs male), and region (bottom, central, peripheral, top), we employed a linear mixed-effects model for differential expression analysis. Initially, single-factor comparisons were performed (e.g., aged vs young, female vs male, region vs region) by pooling across the remaining variables. Subsequently, multi-factor pairwise comparisons were conducted between 16 individual sample groups, isolating the effect of only one controlled variable at a time (e.g., AFB vs YFB, comparing aged female bottom vs young female bottom samples).

### Lung aging is associated with coordinated ECM, metabolic, and immune remodeling

We first examined age-associated differences in the lung proteome. Principal component analysis (PCA) revealed clear separation between aged and young samples along PC1, indicating age as a dominant source of proteomic variance (**Fig. 2a**). A total of 208 proteins with adjusted *p* values (FDR < 0.05) and >1.5-fold changes were considered significantly different between aged and young samples, including 87 upregulated in aged and 121 upregulated in young tissues (**Fig. 2b, Table S5**). Among them, 12 matrisome proteins were elevated in aged samples and 23 in young samples (**Fig. 2b)** with differentially expressed proteins (DEPs) corresponding to rows in the heatmap (**Fig. 2c**). Some upregulated matrisome proteins in aged samples include serine protease inhibitor A3N (SERPINA3N) and fibroblast growth factor 2 (FGF2). SERPINA3N is a protective serine protease inhibitor involved in immune activation and tissue remodeling, while FGF2 promotes fibroblast proliferation and angiogenic signaling, both reflecting active matrix and repair responses in aged lungs.^23,24^ Some upregulated matrisome proteins in young samples include collagen structural components (COL1A1, COL3A1, COL5A1) and collagen crosslink processing enzymes (LOX, LOXL2, PLOD1, PLOD2), representing an anabolic matrix assembly program characteristic of younger lungs.^25^

**Fig. 2:**
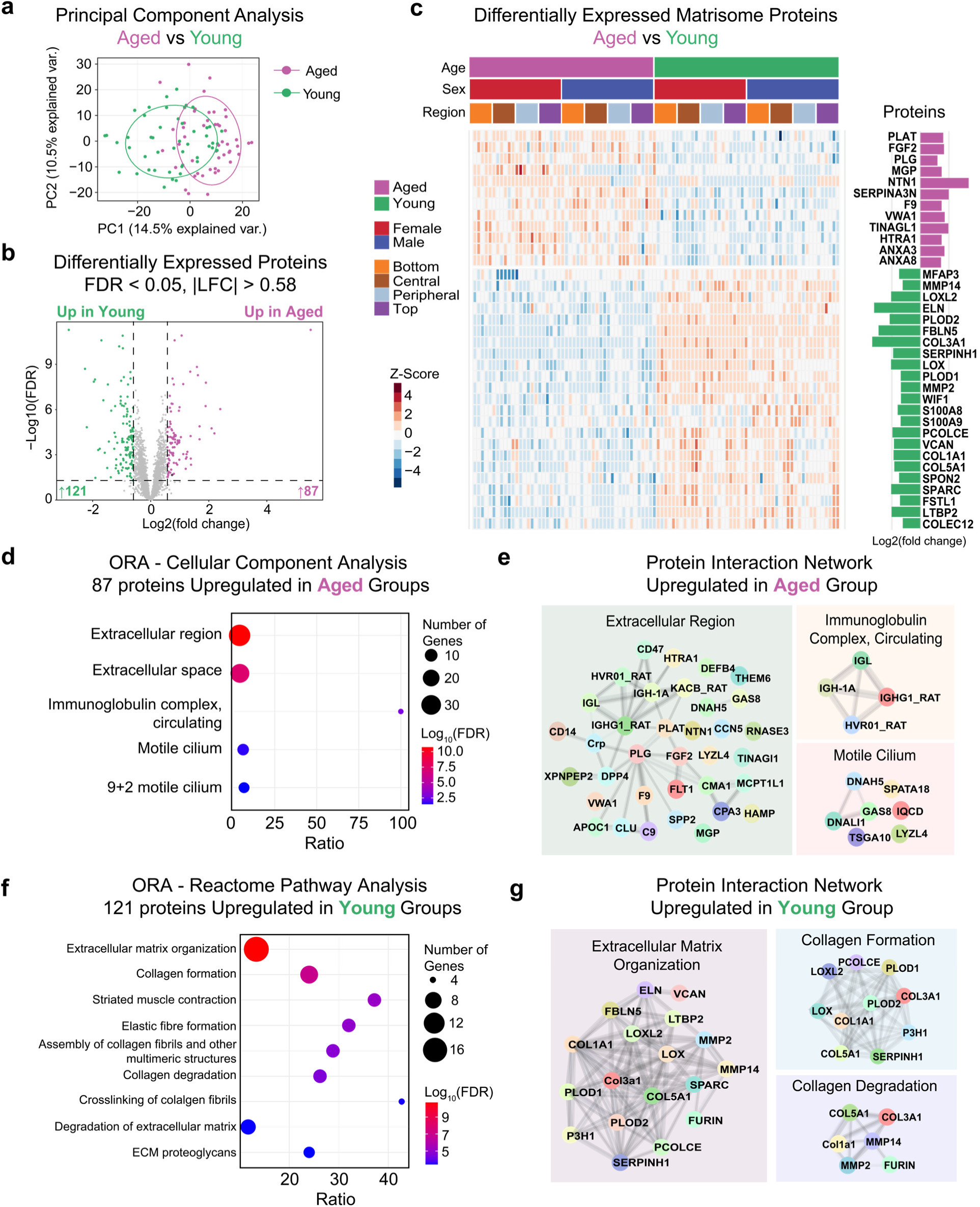
Comparison of aged and young rat lungs reveals distinct proteomic signatures and extracellular matrix remodeling. **a** Principal component analysis of all quantified proteins (n = 3600) comparing aged (magenta) and young (green) rat lung samples. **b** Volcano plot displaying differentially expressed proteins (DEPs) between aged and young groups. **c** Heatmap of differentially expressed matrisome proteins identified from **b**, grouped by age, sex, and region. Each column represents an individual sample (n = 96) and each row a protein. Scaled Z-scores indicate relative expression abundance. Bar on the left indicates Log_2_(fold change). **d** Gene Ontology (GO) over-representation analysis (ORA) of cellular-component terms enriched among proteins upregulated in aged tissues (FDR < 0.05). **e** Protein–protein interaction network of aged-upregulated proteins highlighting clusters corresponding to the extracellular region, immunoglobulin complex, and motile cilium functional groups. Node color reflects subcellular annotation from GO cellular-component categories. **f** Reactome pathway enrichment of proteins upregulated in young tissues showing significant enrichment (FDR < 0.05). **g** Protein–protein interaction network of young-upregulated proteins grouped by major Reactome clusters, including extracellular matrix organization, collagen formation, and collagen degradation.

To identify enriched biological pathways, we performed over-representation analysis (ORA) of the DEPs against Gene Ontology (GO) categories. The 87 upregulated DEPs in aged samples are significantly enriched (FDR < 0.05) in biological processes related to immune response, complement activation and response to bacterium and external stimulus; in cellular components related to extracellular region, extracellular space, immunoglobulin complex and cilium; and in molecular functions related to peptidase activity (**Fig. 2d, Fig. S3a,** and **Table S6**). No Reactome pathways reached significance (FDR < 0.05). Protein interaction network maps generated by STRINGdb highlight the enriched protein hits within GO cellular component clusters (**Fig. 2e**). The extracellular region cluster delineates an extracellular remodeling program that integrates proteolysis, signaling, and matrix-cell interface process, rather than uniform matrix accumulation. Within the network, tissue-type plasminogen activator (PLAT) and plasminogen (PLG) mediate fibrinolysis while serine protease HTRA1 (HTRA1), tubulointerstitial nephritis antigen-like protein (TINAGl1), and von Willebrand factor A domain-containing protein 1 (VWA1) control basement-membrane turnover and adhesion. Matrix Gla protein (MGP) and netrin-1 (NTN1) participate in mineralization and guidance signaling, FGF2 acts as a signaling hub linking fibroblast activation and angiogenesis, and coagulation factor IX (F9) is involved in coagulation processes. These functional roles are broadly established across ECM biology and wound-remodeling literature.^26,27^ Together, these matrisome proteins define a functional remodeling axis characteristic of the aged extracellular environment.

Beyond matrisome proteins, aged-upregulated proteins showed strong enrichment for proteins involved in iron-handling and oxidative-stress pathways, including ferritin light chain 1 (FTL1), ferritin heavy chain (FTH1), and ceruloplasmin (CP). Their coordinated increase suggests enhanced iron sequestration and ferroxidase activity to buffer redox-active iron, consistent with previous reports of ferritin and ceruloplasmin elevation during aging and oxidative imbalance.^28,29^ Concurrently, enrichment of complement C3 (C3), complement C4b (C4B), complement C1q subcomponent subunit A (C1QA), serum amyloid A-1 protein (SAA1), haptoglobin (HP), and beta-2-microglobulin (B2M) suggests activation of both innate and adaptive immune surveillance, reflecting a pro-inflammatory and immune-enriched microenvironment. Collectively, these findings support that the aged lung proteome is defined by the convergence of extracellular remodeling, iron dyshomeostasis, and chronic immune activation, establishing an interconnected ECM–metabolic–immune axis as a hallmark of the aging lung.

ORA against Reactome and GO pathway terms showed that the 121 proteins upregulated in young tissues were strongly enriched for extracellular structure organization and related pathways, including ECM organization and degradation, collagen formation and breakdown, elastin fiber assembly, and ECM proteoglycans (Reactome in **Fig. 2f**; GO terms in **Fig. S3b;** detailed list in **Table S7**). Additional enriched categories such as striated muscle contraction and muscle system development suggest maintained contractile and mechanical integrity in young lungs. The corresponding GO terms reinforced this interpretation, highlighting extracellular matrix, collagen-containing matrix, contractile muscle fiber and actin-binding as dominant cellular component and molecular function features (**Fig. S3b**).

Protein interaction network maps visualize the enriched protein hits in ECM- and collagen-related Reactome clusters (**Fig. 2g**). Representative young-upregulated proteins include COL1A1, COL3A1, COL5A1, elastin (ELN), and fibulin-5 (FBLN5), which are structural constituents of the fibrillar and elastic matrix that preserve tissue elasticity and tensile strength in the lungs;^30,31^ Lysyl oxidases (LOX, LOXL2), together with PLOD1 and PLOD2,, catalyze sequential post-translational crosslink modifications of collagen, hydroxylating and oxidatively deaminating lysine residues to enable glycosylation and covalent crosslink formation which stabilize collagen and elastin fibers;^32^ Matrix metalloproteinases MMP2 and MMP14, which mediate ECM turnover, were also enriched.^33^ The concurrent enrichment of anabolic and catabolic collagen pathways (**Fig. 2f, g**) underscores a dynamic remodeling state that maintains matrix integrity while supporting growth and repair. In sum, young tissues exhibit a coordinated remodeling program dominated by matrix synthesis, fiber assembly, and protease-balanced degradation, defining a structurally adaptable and mechanically resilient younger extracellular environment.

Moreover, Gene Set Enrichment Analysis (GSEA) provided pathway-level insights complementary to the discrete enrichment identified by ORA. While ORA detects over-represented categories among significantly altered proteins, GSEA captures coordinated trends across the entire ranked proteome, unveiling subtle but biologically coherent trends. In the aged group (**Fig. S4a, Table S8**), GSEA confirmed enrichment of reactive nitrogen species metabolism, coagulation, and motile cilium terms, pointing to oxidative and vascular stress responses and epithelial remodeling. In the young group (**Fig. S4b, Table S9**), GSEA demonstrated strong enrichment for ribonucleoprotein complex biogenesis with active transcriptional and translational regulation, which is consistent with robust biosynthetic and regenerative capacity in younger lungs.

### Integrative multi-factor analysis reveals context-dependent aging signatures across sex and region

To study how age, sex, and anatomical region interact to shape the global proteomic landscape, we conducted an integrative multi-factor analysis encompassing all combinations of these variables. Building on the single-factor aged versus young comparison, we next assessed how age-associated differences vary across sexes and lung regions. This multi-factor framework enabled us to disentangle shared and context-specific signatures of aging, extracting both conserved core remodeling programs and sex- or region-selective molecular features that were masked in single-factor analyses.

Across eight pairwise aged versus young comparisons spanning sex and region factors, variable numbers of DEPs were observed, indicating that the magnitude of age-associated remodeling is context-dependent (**Fig. 3a, Table S10**). Female tissues consistently exhibited a greater number of DEPs than male tissues, suggesting higher age sensitivity in the female lung proteome. Regionally, the central and top areas showed more pronounced age-related differences than the bottom and peripheral areas, reflecting spatial heterogeneity. A similar pattern was observed when comparing matrisome DEPs (**Fig. 3b**).

**Fig. 3:**
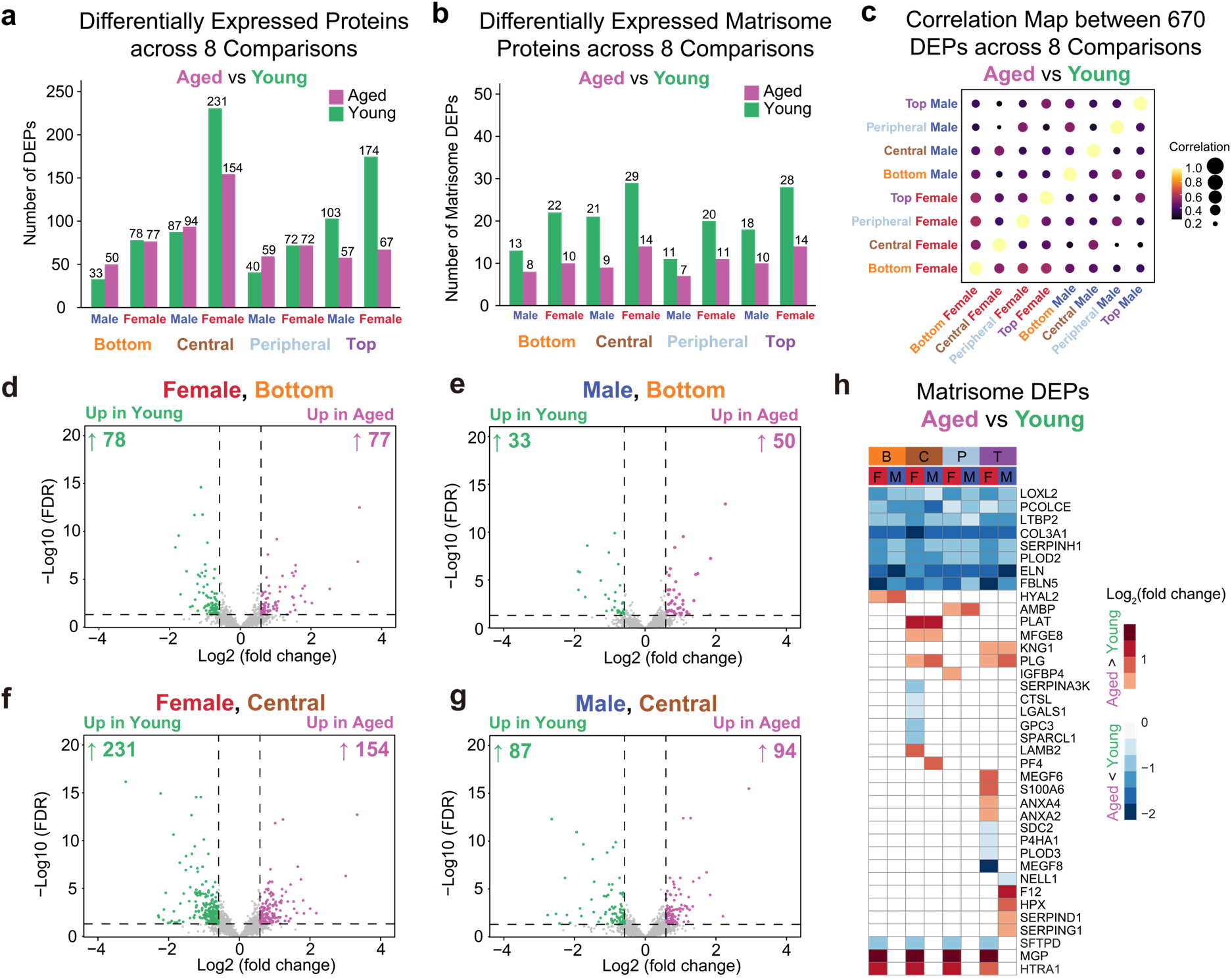
Multi-factor comparisons reveal context-dependent age-associated remodeling across sex and lung regions. **a** Total number of differentially expressed proteins (DEPs) identified in eight pairwise aged versus young comparisons considering sex and region as fixed factors (female and male; bottom, central, peripheral, and top regions). Each pair of adjacent bars represents one comparison, and bar color corresponds to the number of upregulated proteins in aged (magenta) and young (green) tissues. **b** Number of differentially expressed matrisome proteins identified in the same eight comparisons. **c** Pairwise Pearson correlation matrix of differential expression across all eight aged versus young comparisons. **d-g** Volcano plots showing DEPs between aged and young lungs in the (d) female bottom; (e) male bottom; (f) female central and (g) male central groups. Each point represents one protein. Vertical and horizontal lines indicate the cutoffs (fold change = 1.5 and false discovery rate = 0.05). **h** Heatmap showing Log_2_(fold changes) of matrisome proteins significantly altered in at least one of the eight aged versus young comparisons. Blank cells represent proteins that do not pass the filtering threshold or without significant differential expression.

To quantify the coherence of age-associated changes across sexes and anatomical regions, we computed pairwise Pearson correlations among 670 proteins that were differentially expressed in at least one of the eight aged vs young comparisons (**Fig. 3c, Table S11**). The resulting correlation matrix revealed modest positive correlations (r = 0.23 -0.58; median 0.42). The strongest correlations occurred among female regions (r = 0.38 - 0.58), whereas male regions exhibited slightly weaker similarity (r = 0.30 - 0.52). Cross-sex correlations were lower overall (r = 0.23 - 0.49), except when comparing the same region between sexes (**Fig. 3c**). Collectively, these results demonstrate that age-associated protein changes are directionally conserved but display moderate heterogeneity across anatomical regions and sexes, reflecting both shared aging signatures and localized adaptations.

Volcano plots comparing aged vs young proteomes in female and male samples from bottom and central regions, as well as top and peripheral regions, are shown in **Fig. 3d-g** and **Fig. S5a-d,** respectively. Across all eight comparisons, 21 proteins were consistently differentially expressed (points outlined in black in **Fig. 3d-g** and **Fig. S5a-d; also listed in Table S12**), including 10 proteins upregulated in aged and 11 upregulated in young tissues, 8 of which are matrisome proteins. Consistent with the single-factor analysis, aged-upregulated proteins, including FTL1, FTH1, hepcidin (HAMP), clusterin (CLU), vascular endothelial growth factor receptor 1 (FLT1), and secreted phosphoprotein 24 (SPP2), represent iron-handling and stress-adaptive processes, whereas young-upregulated proteins, including LOXL2, PLOD2, FBLN5, ELN, COL3A1, latent-transforming growth factor beta-binding protein 2 (LTBP2), and procollagen C-endopeptidase enhancer 1 (PCOLCE), reflect anabolic and structural ECM maintenance pathways (**Fig. 2c-g**). The uniform directionality of these changes across all comparisons suggests robust and reproducible aging-youth signatures shared across sexes and lung regions.

Besides this conserved DEP set, the heatmap illustrates the Log2 fold change of matrisome proteins showing unique age-dependent expression differences (|fold change| > 1.5, FDR < 0.05) within specific sex and regional combinations (**Fig. 3h**). Proteins without significant changes or proteins which fail to pass the second round of filtering are shown as blank (see Methods). While the overall direction of ECM remodeling was shared across conditions, the magnitude and composition of DEPs varied by both sex and anatomical region. For instance, the first eight matrisome proteins were consistently upregulated in young groups across all comparisons, yet COL3A1 showed particularly high expression in female central regions. Similarly, ELN expression was preferentially higher in male top and bottom regions (**Fig. 3h, Fig. S6a, b**). In contrast, laminin subunit beta-2 (LAMB2) exhibited selective upregulation only in aged female central samples (**Fig. S6c**), suggesting reinforcement of basement membrane in response to mechanical and oxidative stress accumulation. Similarly, annexin A4 (ANXA4) was upregulated only in aged female top samples (**Fig. S6d**), suggesting region-specific epithelial adaptation and membrane repair activity. In sum, these analyses indicate that, although aging elicits a shared global proteomic remodeling trend, its expression pattern is fine-tuned by sex and regional microenvironment, giving rise to distinct molecular trajectories across the lung tissue, which will be discussed in the following sex and region sections.

### Lung aging is associated with changes in collagen and elastin crosslink patterns

To further functionally validate the age-associated ECM remodeling observed in the global proteomic analysis, we quantified major collagen and elastin crosslinks using a targeted LC-MS/MS method optimized for lung tissue. Collagen contains both bivalent immature and trivalent mature crosslinks. Immature bivalent crosslinks include lysinonorleucine (LNL), hydroxylysinonorleucine (HLNL), and dihydroxylysinonorleucine (DHLNL) (**Fig. 4b**). Immature crosslinks can go through additional reactions to form mature trivalent collagen crosslinks: pyridinoline (Pyr) and deoxypyridinoline (DPr) (**Fig. 4b**). On elastin, tetravalent crosslinks, desmosine (Des) and isodesmosine (IsoDes), are formed (**Fig. 4b**). Lung tissues (∼20 mg) from young and aged rats (n = 10/group) were homogenized, chemically reduced, hydrolyzed, and analyzed by triple–quadrupole mass spectrometry (**Fig. 4a**). A representative chromatogram demonstrates baseline separation and detection of all seven crosslink species, with Des and IsoDes co-detected under high-signal conditions following 10-fold dilution (**Fig. 4c**).

**Fig. 4:**
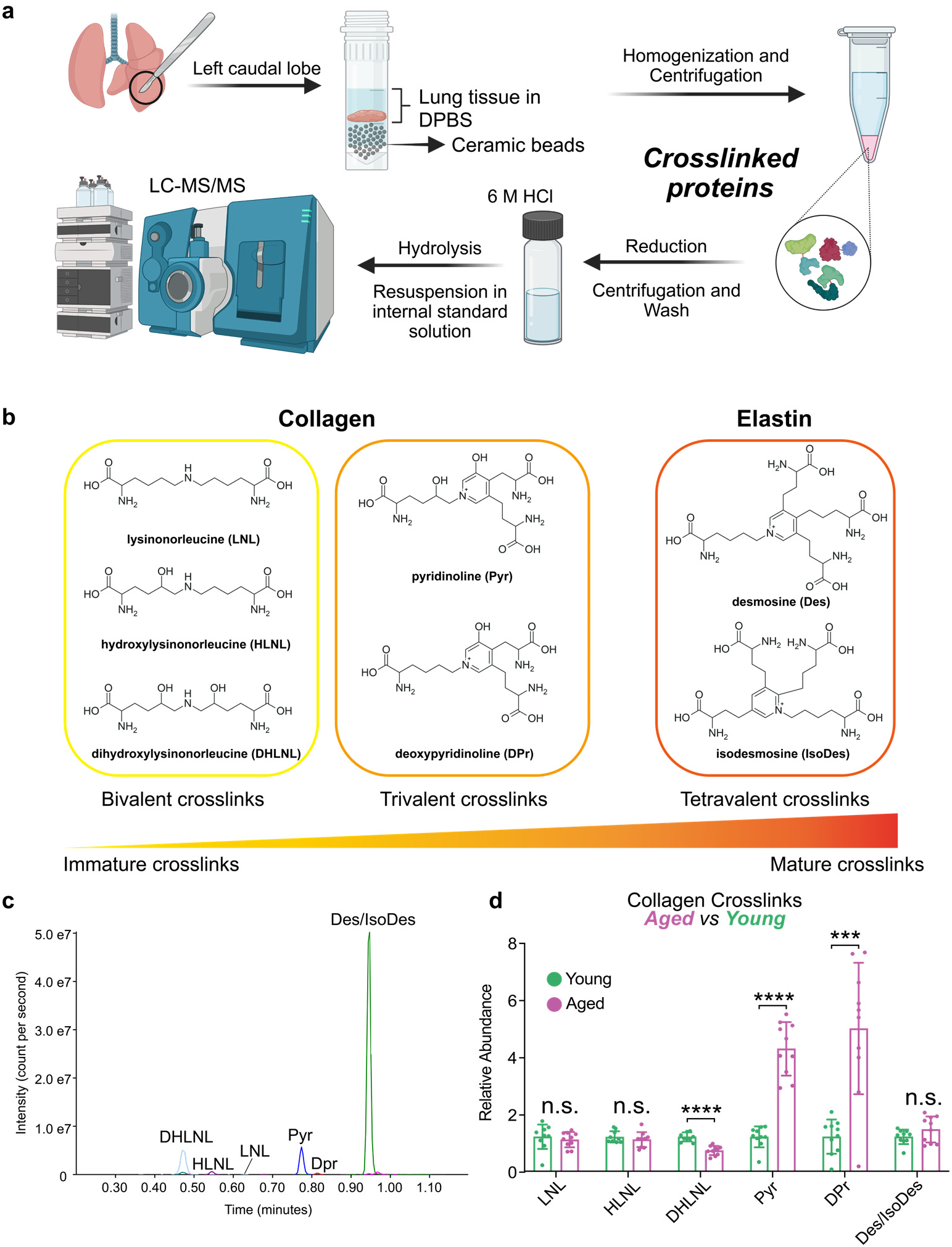
Targeted LC-MS/MS quantification of collagen and elastin crosslinks in young and aged rat lungs. **a** Schematic workflow for crosslink extraction and analysis. Flash-frozen lung tissues were homogenized in DPBS, centrifuged, and the insoluble pellet containing crosslinked ECM proteins was reduced with NaBH₄. Samples were then washed, hydrolyzed in 6 M HCl overnight, and analyzed by LC-MS/MS using internal standards and analyte-specific transitions. **b** Chemical structures of the seven crosslinks categorized by crosslink class and maturity: bivalent (lysino-, hydroxylysino-, and dihydroxylysinonorleucine; LNL, HLNL, DHLNL), trivalent (pyridinoline and deoxypyridinoline; Pyr, DPr), and tetravalent (desmosine and isodesmosine; Des, IsoDes). Crosslinks are arranged on a spectrum from immature to mature, based on known biosynthetic pathways. **c** Representative chromatogram showing baseline separation and detection of seven crosslinks. Peaks are labeled by analyte name. **d** Quantification of individual crosslinks. Relative abundance was determined by analyte-to-internal standard peak area ratio and normalized to tissue weight. Each point represents one biological replicate. Bars indicate mean ± S.D.. Statistical comparisons used two-tailed unpaired Welch’s t-tests: *n.s.,* not significant; ***, *p* < 0.001; ****, *p* < 0.0001.

We reported normalized relative abundance values comparing individual crosslinks in lung tissue from young and aged rats (**Fig. 4d**). Among the immature bivalent crosslinks, LNL and HLNL showed no significant difference between groups, while DHLNL was significantly reduced in aged lungs, suggesting reduced availability of precursors for collagen maturation. This is concordant with reduced expression of immature collagen crosslink enzymes such as LOX, LOXL2, PLOD1 and PLOD2 in aged lungs (**Fig. 2c**). In contrast, the mature trivalent crosslinks Pyr and DPr were markedly elevated in aged samples, indicating increased accumulation of stabilized collagen crosslinks with age. The abundance of Des/IsoDes, elastin-derived tetravalent crosslinks, showed a modest upward trend but did not reach statistical significance. Together, these targeted measurements confirm that aged lung tissue is enriched for mature crosslinks and stable ECM structures, while young tissue exhibits relatively higher immature crosslinks for active fibrogenesis, complementing the global proteomic signature of altered ECM dynamics.

### Sex-dependent proteomic variation and modulation of aging-associated remodeling

Next, we examined the effect of sex on the global lung proteome. PCA indicated largely overlapping distributions of female and male samples, indicating that sex contributes minimally to total proteomic variance compared with age or region factors (**Fig. 5a**). Consistent with this observation, only 16 DEPs were identified between sexes: 10 upregulated in females and 6 upregulated in males (**Fig. S7a, Table S13**). Among 10 female-upregulated proteins, three are matrisome proteins (KNG1, SERPINA6, and MEGF8), suggesting subtle sex-linked variation in extracellular organization and protease regulation (**Fig. S7a**). Kininogen-1 (KNG1) and corticosteroid-binding globulin (SERPINA6), both secreted plasma proteins, participate in protease inhibition and vascular homeostasis. Multiple epidermal growth factor-like domains 8 (MEGF8) contributes to matrix cell signaling and morphogenetic processes. Due to the limited number of DEPs, ORA could not be performed reliably. Instead, we applied GSEA using the complete ranked protein list and identified Reactome pathways significantly enriched in platelet degranulation and Ca^2+^ dynamics (**Fig. S7b**). No significant enrichment was observed against GO items.

**Fig. 5:**
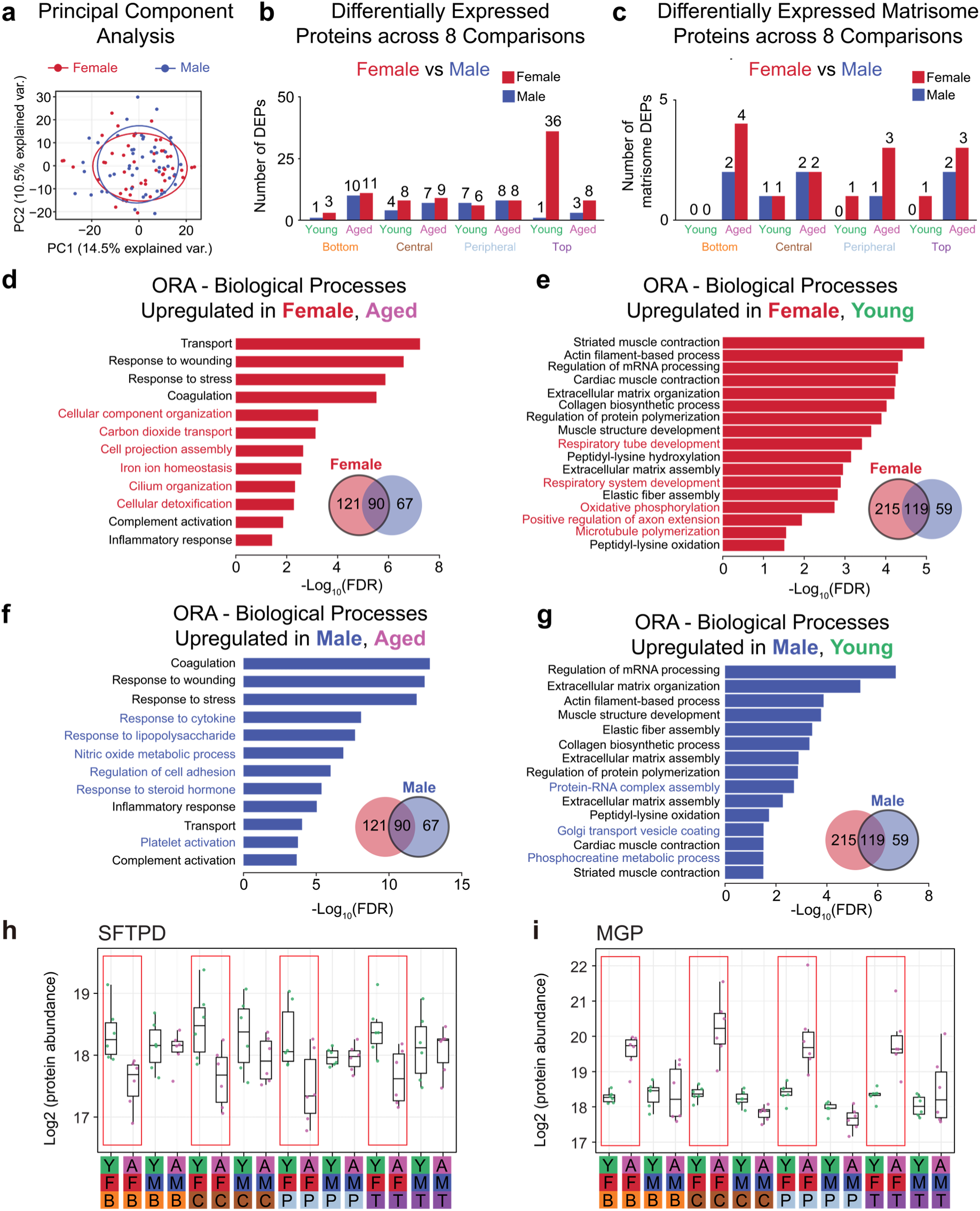
Sex influences the lung proteome and modulates aging-associated remodeling. **a** Principal component analysis of all quantified proteins comparing females and males. **b** Number of differentially expressed proteins (DEPs) identified in eight pairwise female versus male comparisons considering age and region as fixed factors (aged and young; bottom, central, peripheral, and top regions). Each pair of adjacent bars represents one comparison, and bar color corresponds to the number of upregulated proteins in female (red) and male (blue) tissues. **c** Number of differentially expressed matrisome proteins identified in eight pairwise female versus male comparisons. **d-g** Over-representation analysis (ORA) of DEPs in aged versus young from female or male groups. Upregulated proteins used for ORA input are highlighted in black circles from Venn diagrams. Bars represent significantly enriched GO Biological Process terms (FDR < 0.05), with pathway labels color-coded by sex. Pathways shared with single-factor aged versus young comparisons are shown in black, while sex-specific pathways are highlighted in color. **h-i** Example boxplots of two matrisome proteins, SFTPD and MGP, exhibiting sex-specific differences in aged versus young expression. Each point represents one biological replicate.

To assess how sex effects vary with age and lung region, we compared female and male proteomes within each region separately for young and aged groups (**Fig. 5b-c, Table S14**). Overall, the number of sex-associated DEPs remained low across all comparisons, confirming that sex is a minor but context-dependent source of proteomic variation. The largest divergence was observed in the young top region (**Table S15**), where 36 proteins were upregulated in female tissues, enriching for mitochondrial metabolism, muscle contraction and system process by ORA (**Fig. S8, Table S16**). When restricted to the matrisome subset, only 1 to 4 proteins per comparison differed between sexes, indicating limited sex influence on ECM composition at the proteomic level (**Fig. 5c**). These findings demonstrate that sex-linked differences in the lung proteome are subtle overall, with only modest regional and age variation.

Given these minimal baseline sex differences, we next asked whether sex affects age-associated proteomic remodeling. To address this, we reused the DEPs from eight pairwise aged vs young comparisons within each sex across all four regions (**Fig. 3a**). Venn diagrams (**Fig. S9**) and UpSet plots (**Fig. S10**) showed that while a substantial subset of age-associated proteins was shared between sexes, each also displayed distinct DEPs unique to females or males (**Table S17**). Two representative proteins, surfactant protein D (SFTPD) and MGP (**Fig. 5h-i),** show consistent sex divergence across all regions. For example, SFTPD, an innate immune and surfactant-associated protein secreted by alveolar type II cells, was preferentially upregulated in young females (**Fig. 5h**). Similarly, MGP, a calcification inhibitor and vascular matrix regulator, was upregulated in aged females but not males (**Fig. 5i**), reflecting sex-linked differences in proteome expression in both young and aged lungs.

To determine whether these protein-level patterns reflect broader functional trends, we performed ORA using sex-specific aged versus young DEPs and identified both shared and sex-specific biological processes (**Fig. 5d-g, Table S18**). In both aged females and males, upregulated proteins were enriched in processes linked to complement activation, wound and stress adaption coagulation, and inflammatory response (black items) (**Fig. 5d,f**). These items were also enriched in the single-factor aging analysis (**Fig. S3a and S4a**), defining a core aging signature involving immune activation and tissue-repair processes that is broadly conserved between sexes. Beyond this common framework, aged females showed additional enrichment in cellular detoxification, carbon dioxide transport, iron ion homeostasis, and cilium organization, suggesting broader activation of metabolic and oxidative defense mechanisms (**Fig. 5d**). In contrast, aged males exhibited selective enrichment in response to cytokine and lipopolysaccharide, nitric oxide metabolism, and regulation of cell adhesion, reflecting a more focused immune and endothelial remodeling response (**Fig. 5f**). Together, these results suggest that while males and females share a conserved aging framework centered on inflammation and hemostasis, females engage a wider set of adaptive and metabolic pathways, consistent with more extensive tissue remodeling and stress compensation during aging.

In the young-upregulated proteome, both sexes showed enrichment in categories related to extracellular matrix organization, collagen and elastic fiber assembly, muscle structure development, and peptidyl-lysine hydroxylation (**Fig. 5e, g**). These pathways, also identified in the single-factor analysis (**Fig. S3b, S4b**), represent a core young signature characterized by active ECM synthesis and tissue maintenance. Beyond these shared framework, young females displayed additional enrichment in respiratory tube development and oxidative phosphorylation, pointing to enhanced structural and metabolic capacity that supports airway elasticity and energetic homeostasis (**Fig. 5e**). Young males, meanwhile, showed selective enrichment in Golgi transport, creatine metabolism, and protein-mRNA processing, suggesting sex-specific biases in protein trafficking and transcriptional regulation (**Fig. 5g**). Collectively, these results suggest that while both sexes share a young proteomic profile centered on ECM biosynthesis and structural renewal, females exhibit greater enrichment in energy metabolism and epithelial development, whereas protein biosynthesis and regulatory processes are enriched in males.

### Region-dependent proteomic variation and modulation of aging-associated remodeling

To characterize spatial variation within the lung, we compared proteomic profiles across bottom, central, peripheral and top regions. PCA showed partial separation of samples by region, with the central and top regions clustering distinctly from the bottom and peripheral regions, indicating anatomical heterogeneity in protein expression (**Fig. 6a**). Pairwise differential expression analysis confirmed that central and top regions yielded the largest number of total and matrisome DEPs compared with other bottom and peripheral regions (**Fig. S12, Table S19**). For example, between central and peripheral regions, 84 proteins showed higher expression in the central and 20 in the peripheral lung (**Fig. 6b**). Central-enriched matrisome proteins, including COL1A1, COL1A2, decorin (DCN), and versican core protein (VCAN), point to elevated ECM structural and fibroblast-related components in the parenchymal core, whereas peripheral-enriched proteins such as MEGF8 and NTN1 are associated with cell guidance and morphogenesis. The bottom-top comparison identified 14 proteins higher in the bottom and 30 in the top (**Fig. 6c**), indicating regionally distinct metabolic and contractile specializations observed with pathway enrichment. The remaining four pairwise region comparisons are shown in **Fig. S13**. GSEA highlighted clear functional compartmentalization across lung regions (**Fig. 6d-g, Fig. S14, Table S20**). Proteins elevated in the central versus peripheral region were strongly enriched for muscle system processes, actin filament organization, and mitochondrial respiration, consistent with greater contractile and metabolic activity in the central parenchyma where airway and vascular structures converge (**Fig. 6d**). The accompanying Reactome analysis confirmed enrichment of striated muscle contraction and respiratory electron transport, supporting a dual role in mechanical tension and oxidative metabolism (**Fig. 6e**). No significant pathways were enriched in the peripheral region. A similar pattern was observed along the bottom-top axis, where top-enriched proteins showed nearly identical enrichment in contractile and mitochondrial pathways (**Fig. 6f, g**). No significant pathways were enriched in the bottom region. Together, these patterns are consistent with increased contractile and metabolic activity in regions where airway and vascular structures are more prominent.

**Fig. 6:**
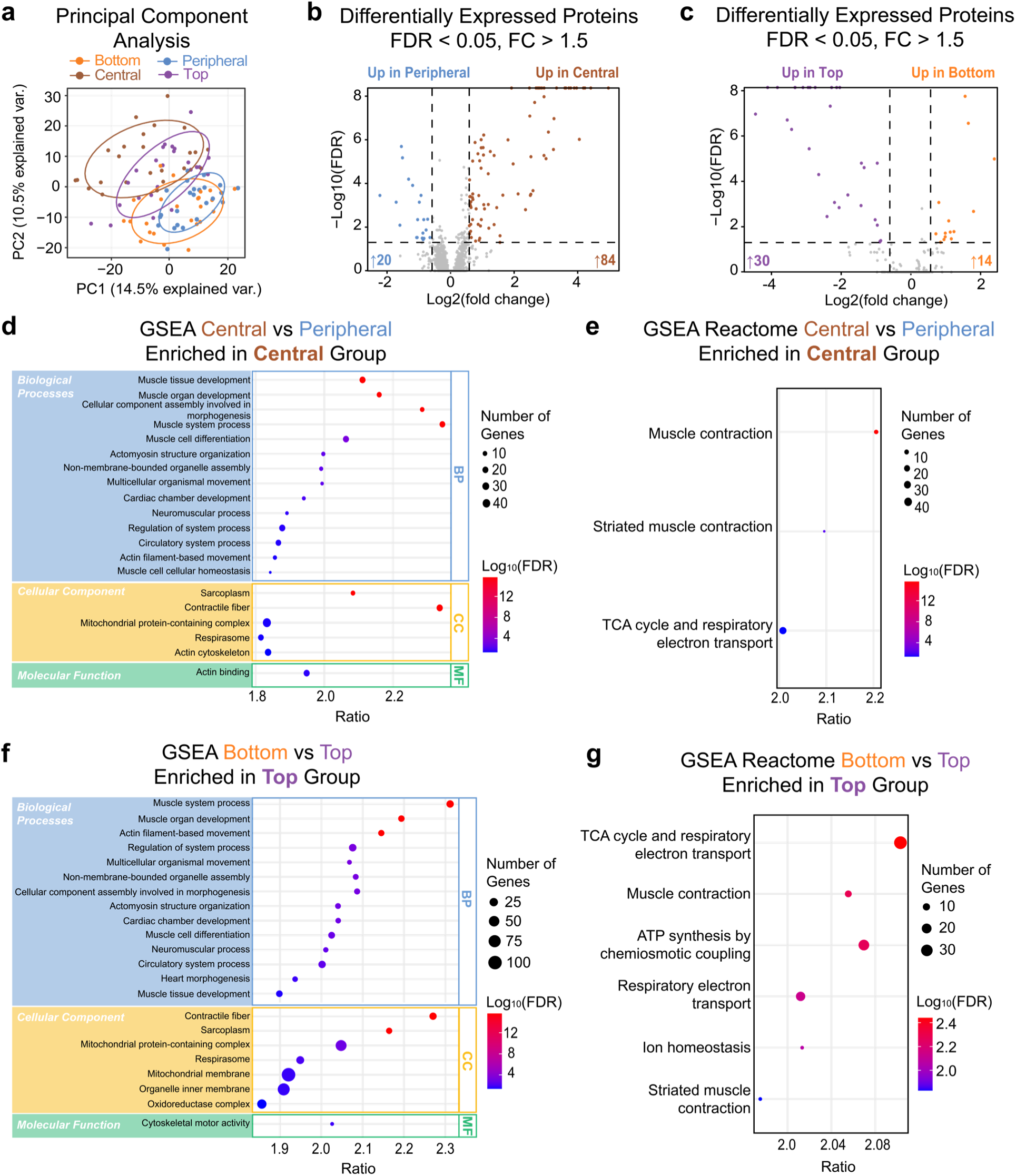
Regional proteomic specialization reveals distinct structural and metabolic programs in the rat lung. **a** Principal component analysis of all quantified proteins comparing anatomical regions. **b** Volcano plot displaying differentially expressed proteins between central and peripheral regions. Proteins with false discovery rate (FDR) < 0.05 and |fold change| > 1.5 are considered significant. Matrisome proteins are highlighted in black circles. **c** Volcano plot comparing bottom and top regions with the same statistical thresholds. **d** Gene set enrichment analysis (GSEA) of the central versus peripheral comparison identifies enriched Gene Ontology Biological Process, Cellular Component, and Molecular Function terms (FDR < 0.05) in the central region. **e** Reactome pathway enrichment of the central versus peripheral comparison in the central region. **f** GSEA of the bottom versus top comparison identifies enriched Gene Ontology Biological Process, Cellular Component, and Molecular Function terms (FDR < 0.05) in the top region. **g** Reactome pathway enrichment of the bottom versus top comparison in the top region. Bubble size indicates the number of overlapping gene hits, and color scale represents –Log₁₀ (FDR).

Building on the single-factor regional analysis, we next examined how regional proteomic differences vary across sex and age. In the young groups, both sexes displayed pronounced regional heterogeneity, with the central-peripheral and central-bottom axes showing the largest number of total and matrisome DEPs (**Fig. S15, and Table S21, S22**). The regional contrast was greater in young females than in young males, and these DEPs are enriched in muscle contraction and system processes with actin binding, reflecting the results of the single-factor analysis (**Fig. S16, Table S23**). These findings suggest that regional specialization is more prominent during youth, particularly in females, potentially reflecting higher basal structural and metabolic differentiation. Inversely, aged males and females exhibited markedly fewer total and matrisome DEPs across regions (**Fig. S15, Table S24**). This convergence of regional profiles implies that aging leads to proteomic homogenization across lung compartments, consistent with reduced tissue plasticity and spatial specialization in the aging organ. Together, these data show that region-dependent proteomic organization is strongest in young lungs and diminishes with age, and that the extent of regional distinction is modulated by sex.

Having established that lung regions exhibit distinct proteomic specializations, we next investigated how regional context modulates age-related remodeling. The Venn diagrams from multi-factor aged vs young analysis across all four regions showed partial overlap among regional DEPs, indicating that while a core set of aging-associated proteins is shared, each region also contributes unique molecular signatures (**Fig. S17**). ORA of aged vs young DEPs in different lung regions showed enrichment in stress response and immune activation in aged tissues (**Fig. S18, Table S25**) while all young regions were enriched in collagen and elastin fiber synthesis, assembly, and extracellular matrix organization (**Fig. S18, Table S25**). These multi-factor results are consistent with the single-factor aged vs young results (**Fig. S3-S4**). Beyond this, the central region displayed the largest number of DEPs when comparing aged and young tissues across the four regions (**Fig. S11, Table S26**), underscoring its role as a metabolically and structurally dynamic compartment sensitive to age-related remodeling. The aged central region showed broad activation of cellular organization and metabolic and immune regulatory pathways, including lipid metabolism and vesicular transport (**Fig. S18b**). The aged peripheral region was characterized by immune-lipid-bacterial response networks, reflecting localized host defense functions (**Fig. S18c**). The aged top region was enriched for nutrient and amino acid metabolism, consistent with its high oxidative demand (**Fig. S18d**), whereas the aged bottom region displayed upregulation of cell migration and organization accompanied with protein degradation, suggesting increased structural turnover (**Fig. S18a**). Conversely, young-upregulated proteins reflected regionally distinct maintenance programs. The central region was dominated by muscle contraction and mitochondrial respiration, consistent with high contractile and metabolic activity (**Fig. S19b**). The top region exhibited enrichment in mRNA metabolism, gene expression, and cytoskeletal organization, reflecting elevated biosynthetic and structural plasticity **(Fig. S19d**). The bottom and peripheral regions in distal lung zones do not contain specific pathways besides common ECM, collagen, elastin and actin organization, consistent with a more uniform structural maintenance role (**Fig. S19a, c**). Together, these results demonstrate that aging remodels the lung proteome by superimposing a shared immune-stress signature onto region-specific structural and metabolic specialization.

## Discussion

This study provides a high-throughput, spatially resolved view of age, sex, and region-dependent remodeling of the lung proteome with an emphasis on the ECM. Using a high-throughput single-step Azo protein extraction coupled to diaPASEF on the timsTOF Pro platform and data analysis in DIA-NN, we achieved deep, reproducible coverage of both total and matrisome protein across a large cohort of 96 rat lung samples. Crucially, collagen and elastin crosslinks were quantified in the aged versus young comparison, enabling direct integration of the ECM crosslink profile with global proteomic remodeling.

A comprehensive, integrated proteomics analysis reveals the effects of aging, sex, and regional variability on pulmonary ECM composition and investigates how these physiological factors collectively influence the lung proteome. The single-factor differential analysis identified overall differences between aged and young, female and male, and region to region comparisons.

Multi-factor analysis further divided individual sample grouping by combining multiple hierarchal factors and considering factor interactions. For example, by fixing the aged vs young comparison grouped by sex and region, we can investigate how sex and region affect the detailed total and matrisome protein composition during aging. This whole workflow provided an unprecedented platform to delineate the lung proteomic landscape within age, sex, and regional comparisons.

Across age, sex, and regional comparisons, age demonstrated the greatest differences in both total protein and matrisome protein expression. Proteomics analysis of the left lung consistently showed enrichment of ECM organization, immune response, and peptidase activity in aged tissues, whereas young lungs were enriched for extracellular structure organization, collagen and elastin fiber assembly, actin/contractile pathways, and surfactant/cilia programs. Proteins enriched in young lungs included ELN, FBLN5, COL1A1, COL3A1, COL5A1, LOX/LOXL2, PLOD1/2, MMP2, and MMP14, consistent with a balance of ECM assembly and protein turnover. In contrast, aged lungs demonstrated enrichment of complement/coagulation pathways, oxidative and iron homeostasis (including FTL1/FTH1 and CP), immune activation, and broader extracellular remodeling. ORA of differentially expressed proteins and GSEA on ranked proteomes demonstrate discrete ECM modules in young tissues and oxidative and immune processes in aging tissues. Combined with our ECM crosslink analysis, these findings suggest that lung aging reflects a shift from a dynamically regulated extracellular network toward a more stress-and inflammation-coupled and mechanically stabilized matrix environment.

Bulk proteomic and transcriptomic studies of aged rodent lungs frequently report activation of stress, inflammatory, and repair-associated pathways, including ECM–receptor interaction, ferroptosis, mitochondrial dysfunction, and impaired regenerative responses, particularly in fibrosis-prone or late-age models.^34–36^ In humans, recent comprehensive proteome atlases provide a global context for aging-associated proteostasis decline and systemic tissue trajectories but are not designed to resolve sex- or region-specific lung architecture or ECM-enriched biochemical fractions.^37^ A prior lung-focused multi-omic analysis further identified age-associated transcriptional programs enriched for extracellular matrix organization and immunoregulatory interactions, alongside reduced metabolic and proliferative pathways, consistent with several aspects of our findings.^38^ In contrast, another human bulk transcriptomic study reported age-associated increases in a subset of structural matrisome proteins (e.g., COL1A1, COL6A1/2, COL14A1, LUM, LTBP4), supported by orthogonal histology.^12^ Notably, the directionality of ECM abundance changes is highly dependent on the aging window analyzed. Here, the comparison between juvenile (7-week) and adult (9-11 month) rat lungs likely captures waning developmental ECM synthesis and ongoing matrix maturation, rather than age-associated fibrotic deposition.

Our multi-factor analyses showed that the age-associated signature was directionally consistent with the single-factor analysis but context-dependent in magnitude across sex and lung region. Females generally exhibited more age-sensitive DEPs than males. Baseline sex differences were modest yet consistent with biological nuance. For example, SFTPD was higher in young females across all regions, suggesting a decrease in immune–surfactant turnover with age, whereas MGP was higher in aged females, pointing to sex-linked vascular/ECM regulation. Regionally, central/top compartments displayed stronger age effects than bottom/peripheral areas and were enriched for muscle/contractile and mitochondrial respiration pathways, consistent with airway/vascular convergence and higher mechanical load.

Cardiovascular proteins including myosin heavy chains and cardiac troponin were identified and quantified in rat lung tissues. The detection of cardiac proteins in lungs is unique in rats due to the extension of the myocardium into pulmonary veins.^39^ We observed that several cardiovascular-associated proteins, including myosin-3 (MYH3), myosin-6 (MYH6), cardiac troponin I (TNNI3), and cardiac troponin T (TNNT2), showed marked upregulation in central and top regions compared with other lung areas. During central and top region dissection and tissue collection, portions of proximal blood vessels or adjacent cardiac tissue may have been included, potentially explaining the strong expression of these cardiac and vascular proteins in the central samples. Moreover, this enrichment is significantly pronounced in young rats (both in female and male) compared with aged rats (**Table S23, S24**). Therefore, the localized dissection bias alone cannot explain the enrichment of proteins involved in contractile and metabolic processes in the central and top regions (**Fig. 7e, g**). Young rats show regional expression discrepancy explicitly along the central-peripheral and bottom-top axes compared with aged rats. Because central region dissections can include proximal vasculature or adjacent myocardium, we emphasize pathway-level rather than single-marker attribution.

The observed reduction in spatial molecular heterogeneity from young to aged rats should be interpreted with caution and considered in the context of both species-specific biology and experimental scale. Human lung aging is frequently associated with increased molecular and cellular heterogeneity, as revealed by transcriptomic and single-cell studies emphasizing immune activation, senescence, and localized remodeling within specific microenvironments.^40,41^ In contrast, our tissue-level proteomic analysis integrates signals across anatomically defined lung regions in a rat model, capturing aggregate molecular programs rather than cell-type–resolved variation. Differences in spatial resolution and species context may therefore underlie the apparent contrast between increased cellular heterogeneity reported in human studies and the reduced regional proteomic contrast observed here.

Aging was the dominant driver of variance across global proteomics and crosslink chemistry data layers. In the crosslink panel, aged lungs showed a maturation shift of the collagen network: pyridinoline (Pyr) and deoxypyridinoline (DPr) increased, while the immature divalent crosslink DHLNL decreased, suggesting slow turnover instead of active fibrogenesis. Additionally, desmosine/isodesmosine (Des/IsoDes) crosslinks from elastin rose modestly. Mechanistically, even if collagen and elastin crosslink enzymes (LOX and PLOD) decline with age in our proteomic analysis, long-lived collagen accrues enzymatic mature links and non-enzymatic aged crosslinks, both contributing to increasing lung stiffness over time. This is consistent with the crosslink results observed here and in previous aging lung reports.^25,42,43^

Using the same method, we quantified the collagen and elastin crosslink events comparing human lung tissues from IPF patients and healthy control donors. In contrast to the normal aging process, IPF lungs show a robust increase in novel immature DHLNL crosslinks, while other mature crosslink species remain unaltered.^42,44^ This result is also validated in a bleomycin fibrotic mouse model.^42^ Aging is well-documented risk factor for the development of IPF. However, an interesting discrepancy is observed when comparing collagen crosslinks between aging lungs and IPF lungs: aging lungs exhibit a stable and mature collagen crosslink network while IPF lungs initiate new collagen crosslinks. These results carry translational implications. First, they establish a baseline aging signature (↑PYD/DPD, ↓DHLNL; modest Des/IsoDes) that helps distinguish physiology from the pathologic ECM remodeling observed in IPF. Second, they motivate composite biomarker panels combining ECM enzymes (LOX/LOXLs and PLOD1/2/3), structural proteins, and crosslink species (DHLNL, PYD, DPD, Des/IsoDes) to distinguish physiological maturation from active fibrogenesis. Third, they support targeted intervention strategies that modulate crosslink quantity and quality—via LOX/LOXL paralog-aware approaches, PLOD-mediated hydroxylation, and oxygen/iron/copper–redox biology—rather than focusing solely on total collagen mass.^45,46^

This study has several limitations that suggest future research directions. The study compared young (7 weeks) to early–mid aging (9–11 months) rather than advanced senescence. Further studies including mice of more advanced ages which will inform longitudinal aging trajectories. Additionally, collagen crosslink quantitation used a surrogate matrix; Des/IsoDes were not separated from one another, and HLNL lacked a certified standard. Therefore, absolute values are method-referenced, and directionality is the primary read-out. Although two-stage imputation and stringent criteria (≥5 real values of 6 per group) strengthen differential-expression inference, targeted validation of representative sex and region-modulated proteins remains warranted. Expanded UniProtKB rat proteome reference could also boost proteome coverage. Finally, regional dissections in the central compartment may include vascular or cardiac tissue. Emerging spatial omics technologies with subcellular resolution, such as Xenium in situ platforms, will enable refined mapping of cell-type–specific landscapes underlying lung aging across anatomical regions.

In conclusion, this study provides the first comprehensive, mass spectrometry-based proteomics analysis of the pulmonary rat proteome aimed at characterizing the effects of age, sex, and lung region on relative protein expression. In tandem with ECM-crosslinking analysis, the results of this study demonstrate the dominant role of age in the structure and composition of the pulmonary ECM.

## Methods

### Chemicals and Reagents

All chemicals and reagents were purchased from Sigma-Aldrich unless otherwise specified. Pierce C18 Tips for peptide desalting were purchased from Thermo Fisher Scientific.

### Histological Assessment

Right lung lobes from 12 young (7-week old) and 12 aged (9-11-month old) SD rats were collected at necropsy, inflated with formalin, processed to paraffin block, sectioned at 5 μm, and stained with hematoxylin and eosin (H&E). Slides were digitally scanned with a Panoramic P1000 scanner (3DHistech) and qualitatively evaluated by a board-certified veterinary pathologist.

### Extracellular Matrix Protein Extraction from Rat Lung Tissue

The extraction of proteins from rat lung tissue was carried out according to previously reported protocols for the solubilization of ECM proteins from lung tissue enabled by the photocleavable surfactant Azo.^20,21^ For each sample, 30 mg lung tissue was weighed out, cryopulverized in liquid nitrogen, and rinsed in Dulbecco’s phosphate-buffered saline (DPBS) to deplete highly abundant serum proteins which may otherwise prevent the detection of lower abundance proteins. The pellet remaining after two rounds of DPBS washing was solubilized in an Azo extraction buffer according to previously reported methods.^20,21^ Following protein extraction, 50 µg protein was reduced by 25 mM TCEP, alkylated with 30 mM chloroacetamide, and digested with Trypsin Gold (Promega). Peptide de-salting was performed with Pierce C18 Tips (Thermo Scientific), and peptides were re-suspended in 0.1% formic acid in water. NanoDrop concentrations were used to determine LC-MS injection volumes.

### Bottom-up Proteomics Data Acquisition

Liquid chromatography (LC) and mass spectrometry methods for bottom-up proteomics data acquisition followed previously reported parameters.^20,21^ Data acquisition was performed on a Bruker nanoElute nanoflow ultra-high pressure LC system in tandem with a Bruker timsTOF Pro. Approximately 400 ng peptides were injected onto an IonOpticks Aurora Ultimate C18 column (25 cm x 75 µm x 1.7 µm) and separated over a 2-hour gradient consisting of 2-17% mobile phase B (MPB) over 60 minutes, 17-25% MPB between 60 and 90 minutes, 25-37% MPB between 90 and 100 minutes, 37-85% MPB from 100 to 110 minutes, and a 10-minute wash step at 85% MPB (mobile phase A: 0.1% formic acid in water; mobile phase B: 0.1% formic acid in acetonitrile). The timsTOF Pro was operated in data-independent acquisition (DIA) mode with parallel-accumulation serial fragmentation (PASEF) enabled. The mass scan range was 400 m/z to 1700 m/z. 400 ng peptide injections of K562 whole cell lysate were used to verify sample injection quantity by comparing total ion current intensities.

### DIA-NN Protein Identification and Quantification

Raw MS/MS data were searched in DIA-NN version 1.8.1 against a spectral library generated using a high-quality, Swiss-Prot reviewed rat FASTA downloaded from UniProt database (8,202 entries, canonical, downloaded 05 September 2024). The protease was set to Trypsin/P with up to 2 missed cleavages. N-terminal methionine excision and carbamidomethylation were selected as fixed modifications. Variable modifications included methionine oxidation and N-terminal acetylation with a maximum of 2 variable modifications. Mass accuracy was set to 10.0 ppm, MS1 accuracy to 20.0 ppm, and a precursor false discovery rate of 1.0% was used.

### Proteomics Data Processing

Proteomics data processing was performed in the R framework (version 4.2.2). The protein abundance data (report.pg_matrix) (containing 3859 protein groups in 96 sample groups, with missing values) were filtered to ensure that each protein group contained at least > 50% real values in all 96 sample or 5 or 6 real values in any of 16 sample types. After the first round of filtering, 3600 protein groups remained. The protein abundances were then Log2-transformed and normalized using the “medianNormalization” function in the “NormalyzerDE” R package (version 1.16.0).

Remaining missing values were imputed by random forest using the “missForest” function in the “missForest” R package (version 1.4). Additionally, if a protein group had no observed values in one sample group, the corresponding random forest-imputed values were reset to NA and re-imputed using the impute.QRILC function in the imputeLCMD R package (version 2.1). This approach ensured that imputed values in sample groups with no observed signal remained below the lower limit of quantification. In contrast, random forest imputation alone, which assumes missing-at-random behavior, could generate artificially high values in such cases and distort expression directionality in downstream differential expression analysis. For sample groups containing at least one observed value, random forest imputation was retained because it preserved local abundance patterns.

Rat matrisome annotations are assigned based on mouse homological matrisome gene symbol annotations from Matrisome AnalyzeR.^22^

Principal component analysis was performed based on total protein (n =3600) expression abundance after filtering and two rounds of imputation using the “prcomp” function in the built-in R function. A 95% confidence interval was used to generate oval outlines.

### Differential Expression Analysis

Differential expression (DE) analysis was performed using a linear mixed effects model implemented in the “lme4” (version 1.1-34) and “lmerTest” (version 3.1-3) R packages.^47,48^ A single-factor differential expression analysis was performed first without considering factor interactions in the “lmer4” function. We compared aged versus young (48 aged vs 48 young), female versus male (48 female vs 48 male) and region vs region (6 pairwise comparisons from Bottom, Central, Peripheral and Top regions). For example, in the central versus peripheral comparison, all 16 central tissue samples were compared with all 16 peripheral tissue samples, regardless of age and sex. Proteins with Benjamini–Hochberg adjusted p-values (false discovery rate or FDR) less than 0.05 combined with estimate changes (Log2 [fold change]) greater than 0.585 (|fold change| > 1.5) were considered statistically significant.

We next performed a multi-factor differential expression analysis on individual sample groups, considering factor interactions in the “lmer4” function. For example, instead of comparing all 48 aged samples with all 48 young samples, we compared n=6 female, bottom, aged tissues with n=6 female, bottom, young tissues. For aged vs young comparisons, there are 8 pairwise comparisons with fixed sex and region factors: AFB versus YFB, AMB versus YMB, AFT versus YFT, AMT versus YMT, AFC versus YFC, AMC versus YMC, AFP versus YFP, AMP versus YMP (see **Fig. 1** for key). For female versus male comparisons, there are 8 pairwise comparisons with fixed age and region factors: YFB versus YMB, AFB versus AMB, YFC versus YMC, AFC versus AMC, YFP versus YMP, AFP versus AMP, YFT versus YMT, AFT versus AMT. For region comparisons, there are 24 pairwise comparisons with fixed age and sex factors: Young Female group (YFB versus YFC, YFB versus YFP, YFB versus YFT, YFC versus YFP, YFC versus YFT, YFP versus YFT), Young Male group: (YMB versus YMC, YMB versus YMP, YMB versus YMT, YMC versus YMP, YMC versus YMT, YMP versus YMT), Aged Female group (AFB versus AFC, AFB versus AFP, AFB versus AFT, AFC versus AFP, AFC versus AFT, AFP versus AFT), Aged Male group (AMB versus AMC, AMB versus AMP, AMB versus AMT, AMC versus AMP, AMC versus AMT, AMP versus AMT).

In a single pairwise comparison, such as AMB versus YMB, a protein group was retained only if at least one of the two groups contained 5 or 6 observed values out of 6 replicates, thereby avoiding comparisons driven solely by imputed values. After this second round of filtering in each comparison, proteins with Benjamini–Hochberg adjusted p-values (FDR) were calculated based on filtered out protein groups in each comparison. Proteins with FDRs less than 0.05 combined with estimate changes greater than 0.585 (|fold change| > 1.5) were considered statistically significant. Estimate change is similar to Log2(fold change), but these values are not identical when using a linear mixed effects model considering multi-factor interactions. However, Log2(fold change) is used for multi-factor comparison figures to facilitate consistent labeling across both single-factor and multi-factor analyses.

### Heatmap Plot

Differentially expressed matrisome proteins in aged versus young single-factor comparisons (FDR < 0.05, |FC| > 1.5) were extracted as our target list in **Fig. 2c**. The protein expression abundance of the target protein list across all individual samples was used to generate the heatmap. Each row corresponds to one protein group. Each column corresponds to one sample out of 96. These rows were clustered based on the Pearson correlation coefficient of their expression profiles across the samples, grouping proteins with similar expression patterns together. The normalized protein expression is scaled and clustered based on row or protein. The columns are listed based on their original sample grouping and are unclustered.

Differentially expressed matrisome proteins in 8 aged versus young multi-factor comparisons (FDR < 0.05, |FC| > 1.5) were extracted in **Fig. 3h**. If a matrisome protein group demonstrated differential expression in any of the 8 pairwise comparisons, the aged vs young Log2(fold change) is listed in the heatmap matrix. Otherwise, the value is listed as NA or left as a blank space. This indicates that either the protein does not show differential expression (NA) or is filtered out in the second round of filtering (blank space).

### Pathway Analysis

Over-representation analysis (ORA) was performed using differentially expressed proteins (FDR < 0.05, |FC| > 1.5) in either single-factor or multi-factor comparisons, and Stringdb and Panther were used to generate Reactome pathways as well as Gene Ontology (GO) terms including Biological Process, Cellular Component, and Molecular Function for rat. Generally, ORA was performed on input protein set sizes greater than 50 differentially expressed proteins. Panther was used to generate ORA Reactome pathways as well as GO terms of Biological Process, Cellular Component and Molecular Function.^49^ If there were too few differentially expressed proteins in each comparison to identify over-represented pathways or terms, gene set enrichment analysis (GSEA) was performed from the total protein list using the 2024 WebGestalt interface: https://www.webgestalt.org/ (access on July 2025).^50^

### ECM Crosslink Analysis in the Rat Lung

Sample preparation and LC-MS/MS analysis of crosslinks in rat lungs were performed according to a previous report.^42^ Briefly, rat lung tissue (∼20 mg) was homogenized in DPBS using an Omni Bead Mill Homogenizer. The homogenate was centrifuged at 15,000 × *g* for 10 minutes at 4 °C. The given pellet was resuspended in 1 mg/mL NaBH_4_ in 0.1 M NaOH and placed on a shaker for 1 hour at 4 °C. The mixture was then quenched with acetic acid (0.1%) and centrifuged at 15,000 × *g* for 10 minutes at 4 °C. After washing with water three times, the pellet was hydrolyzed in 2 mL of 6 M HCl at 110 °C overnight. The mixture was dried down under nitrogen at 80 °C. The residue was reconstituted in 500 μL of an internal standard solution for LC-MS/MS analysis.

Analysis was performed on a Waters CORTECS C18 column (2.1 × 50 mm, 2.7 μm) using an Agilent 1290 UPLC (Agilent, CA) and PAL autosampler (CTC Analytics AG, Switzerland) system coupled with a SCIEX 6500^+^ triple–quadrupole mass spectrometer. The mobile phases for the analysis were 50 mM HFBA in water (mobile phase A) and acetonitrile (mobile phase B) and were delivered at a flow rate of 1.5 mL/min with a gradient elution (0-0.2 minutes, 8% B; 0.2-1.0 minutes, 8%-28% B; 1.0-1.02 minutes, 28%-98% B; 1.02-1.4 minutes, 98% B; 1.4-1.45 minutes, 98%-8% B). D4-desmosine was used as the internal standard for desmosine/isodesmosine, while d4-dihydroxylysinonorleucine was used as the internal standard for all other analytes. The calibration curve samples were made by mixing all analytes (hydroxylysinonorleucine standard was not available) with a range from 0.3 to 1000 ng/mL. The curves were fitted using a linear or quadratic fit and passed the accuracy (within 20%) and precision (within 15%) criteria. The relative abundance of crosslinks in lung tissue was calculated by the peak area ratios of the analyte versus the internal standard and normalized by tissue weight.

## Supplementary Tables

**Table S1** Detailed sample metadata of all 96 samples including group labeling.

**Table S2** Raw DIA-NN output protein group file

**Table S3** Raw DIA-NN output peptide file

**Table S4** Complete protein expression abundance matrix after filtering and imputation

**Table S5** Aged versus Young single-factor differential expression analysis protein list (|fold change| > 1, FDR < 0.05)

**Table S6** Selected enriched GO and Reactome terms of differentially expressed proteins in aged tissues from aged vs young single-factor comparison

**Table S7** Selected enriched GO and Reactome terms of differentially expressed proteins in young tissues from aged vs young single-factor comparison

**Table S8** Selected enriched pathways in aged tissues from gene set enrichment analysis of protein Log2(fold change) list from aged vs young single-factor comparison

**Table S9** Selected enriched pathways in young tissues from gene set enrichment analysis of protein Log2(fold change) list from aged vs young single-factor comparison

**Table S10** Aged versus Young multi-factor differential expression analysis protein list (|fold change| > 1, FDR < 0.05)

**Table S11** 670 total protein and 38 matrisome protein Log2(fold change) and false discovery rate lists showing differential expression in at least one of eight pairwise multi-factor aged vs young comparison

**Table S12** Overlapping proteins up in aged or young tissues in all eight aged vs young multi- factor comparisons

**Table S13** Female versus male single-factor differential expression analysis protein list (|fold change| > 1, FDR < 0.05)

**Table S14** Female versus male multi-factor differential expression analysis protein list (|fold change| > 1, FDR < 0.05)

**Table S15** Differentially expressed proteins from multi-factor female vs male grouped by age and region

**Table S16** Selected enriched GO and Reactome terms of differentially expressed proteins in female tissues from female vs male multi-factor comparison in top young group.

**Table S17** Differentially expressed proteins from multi-factor aged vs young grouped by sex.

**Table S18** Over-representation analysis of differentially expressed proteins from multi-factor aged vs young grouped by sex.

**Table S19** Region versus region single-factor differential expression analysis protein list (|fold change| > 1, FDR < 0.05).

**Table S20** Enriched pathways in aged tissues from gene set enrichment analysis of protein Log2(fold change) list from region vs region single-factor comparison.

**Table S21** Region versus region multi-factor differential expression analysis protein list.

**Table S22** Region versus region multi-factor differential expression analysis protein list (|FC| > 1, FDR < 0.05).

**Table S23** Selected enriched GO and Reactome terms of differentially expressed proteins in central tissues from central vs bottom or peripheral multi-factor comparisons in young groups (either male or female).

**Table S24** Differentially expressed proteins from region vs region comparison grouped by age and sex.

**Table S25** Over-representation analysis of differentially expressed proteins from multi-factor aged vs young grouped by region.

**Table S26** Differentially expressed proteins from multi-factor aged vs young grouped by region.

## Acknowledgements

AbbVie Inc. provided the funding for this project.

## Data and code availability

The mass spectrometry proteomics data have been deposited to MassIVE repository with the identifier MSV000102094.

The R scripts are available upon reasonable request to corresponding authors. We built an RShiny webpage interface to visualize differential expression with multi-factor combinations on: https://feiwang-outobi.shinyapps.io/DEP-volcano-plot3/ and https://feiwang-outobi.shinyapps.io/Protein-combined-analysis3/.

## Authorship information

### Contributions

A.G.T, F.W, Y.H., Y.T., and Y.G. conceptualized and contributed to study design. W.B. and L.P. performed the histological assessment. A.G.T performed the proteomics experiment with assistance from L.J.B. J.Z. performed ECM crosslinking experiment and data analysis. Q.J. mentored ECM crosslinking experiment and analysis. F.W., Y.B., A.J.P., and J.L. participated in conceptualization of proteomics data processing and analysis. F.W. performed proteomics data processing and statistical analysis. F.W. developed the interactive RShiny data visualization platform. A.G.T. and F.W. prepared figures for publication. Y.Z. and T.J.A. served as resources for proteomics experimental design. K.K and L.M.S. contributed as ECM biology experts. A.G.T., F.W., L.M.S., Y.H., Y.T., and Y.G. contributed to writing, reviewing, and editing the manuscript. Y.H., Y.T., and Y.G. supervised the study.

## Supplementary Figures

**Fig. S1:**
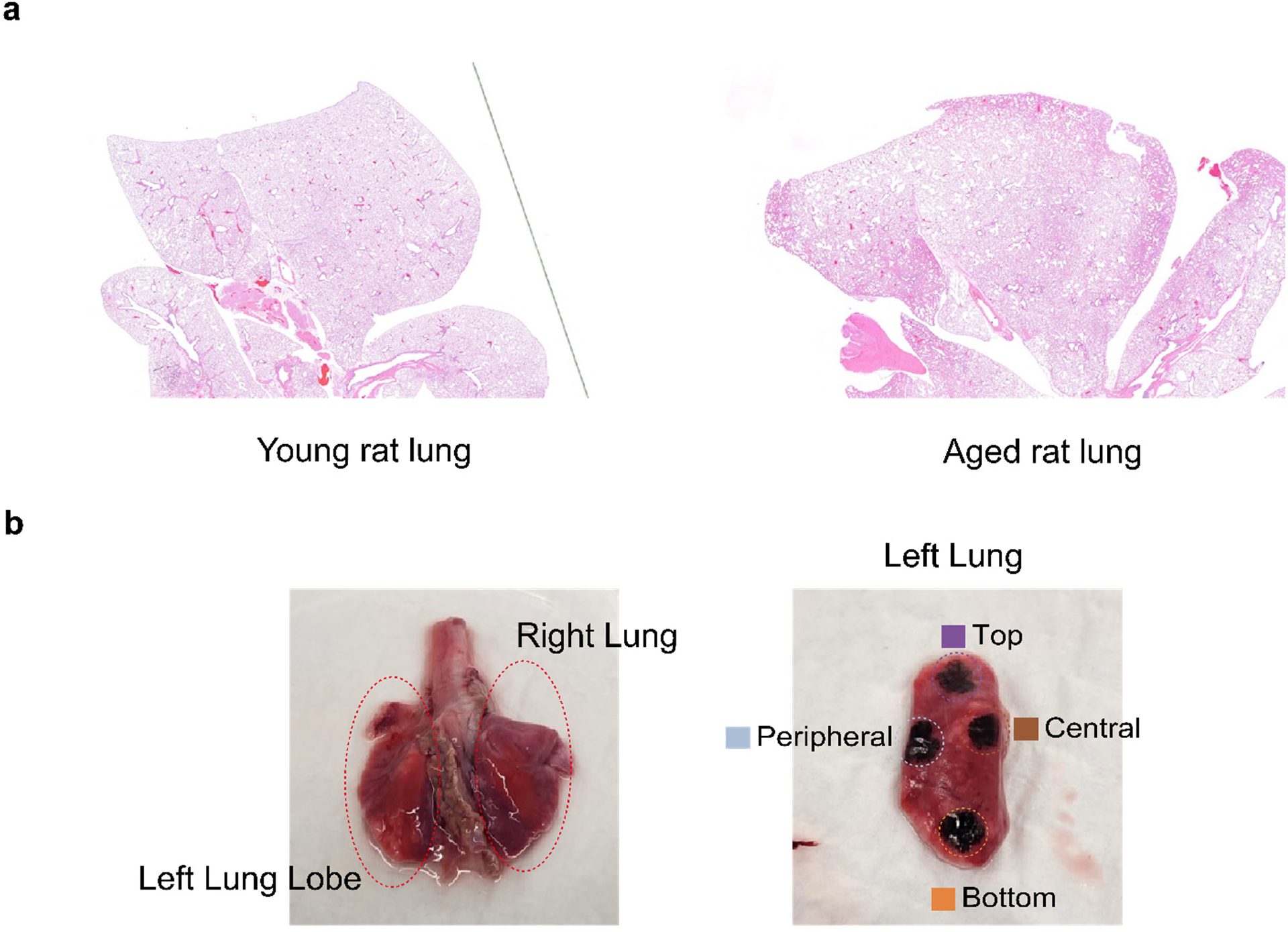
Histological assessment and regional dissection of rat lungs. **a** Representative hematoxylin and eosin (HE) staining of right lungs from young and aged rats. Young lungs showed normal alveolar structure, whereas aged lungs displayed mild macrophage accumulation in subpleural regions. No tumors or major pathological changes were observed. **b** Gross view of lungs and regional sampling scheme. Four regions, bottom (B), central (C), peripheral (P), and top (T), were dissected for proteomic analysis to capture spatial variation.

**Fig. S2:**
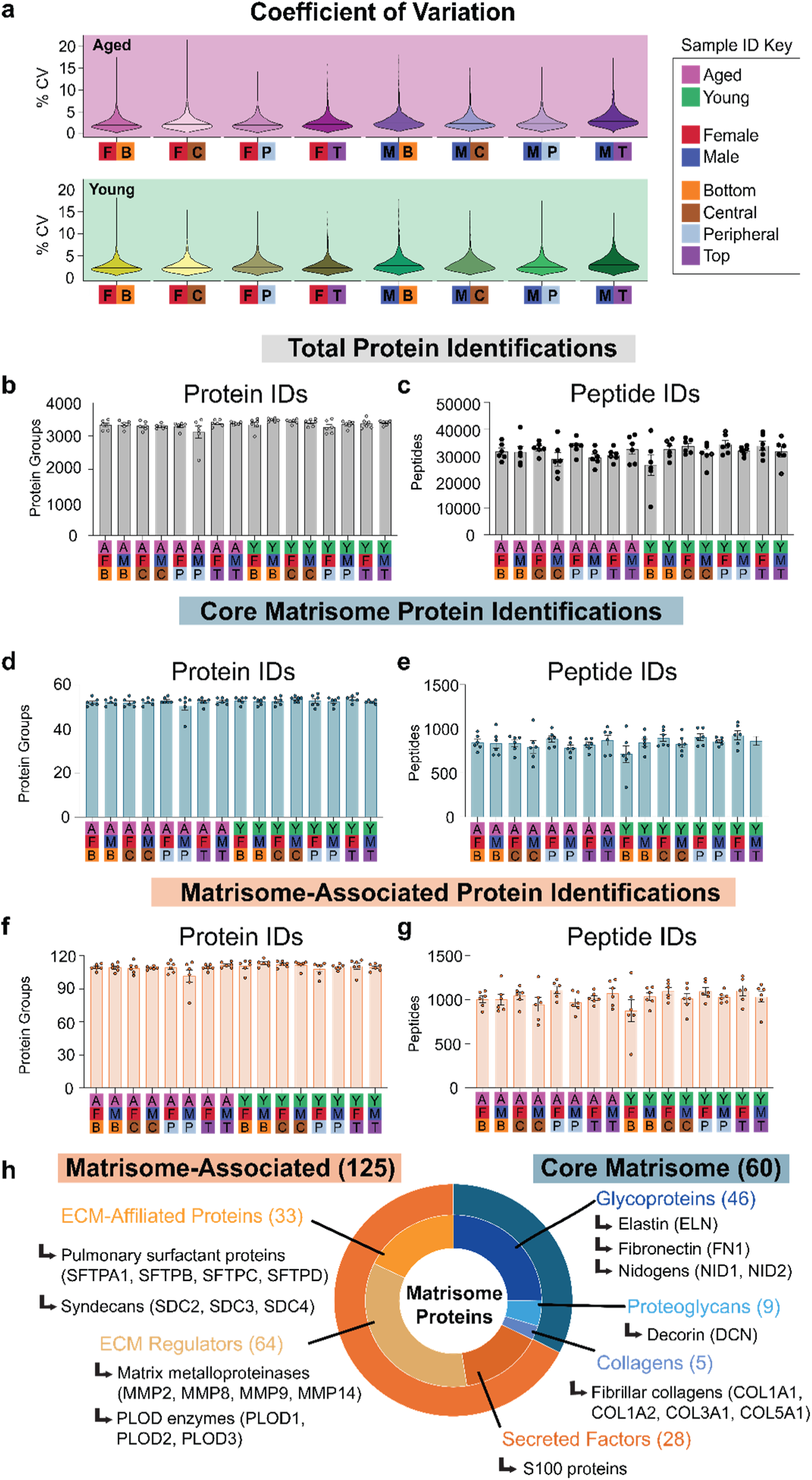
The rat lung proteomics dataset demonstrates high reproducibility and comprehensive proteome coverage. **a** Coefficient of variation (CV) analysis across 16 experimental groups demonstrates high reproducibility of the single-step Azo extraction and diaPASEF workflow. Violin plots show consistent CV distributions across age, sex, and lung regions with most values below 10%, indicating stable quantification. **b, c** Total protein and peptide identifications across all sample groups. **d, e** Core matrisome protein and peptide identifications across all sample groups. **f, g** Matrisome-associated protein and peptide identifications across all sample groups.**h** Summary of the matrisome composition detected in this study, classified into core and matrisome-associated categories.

**Fig. S3:**
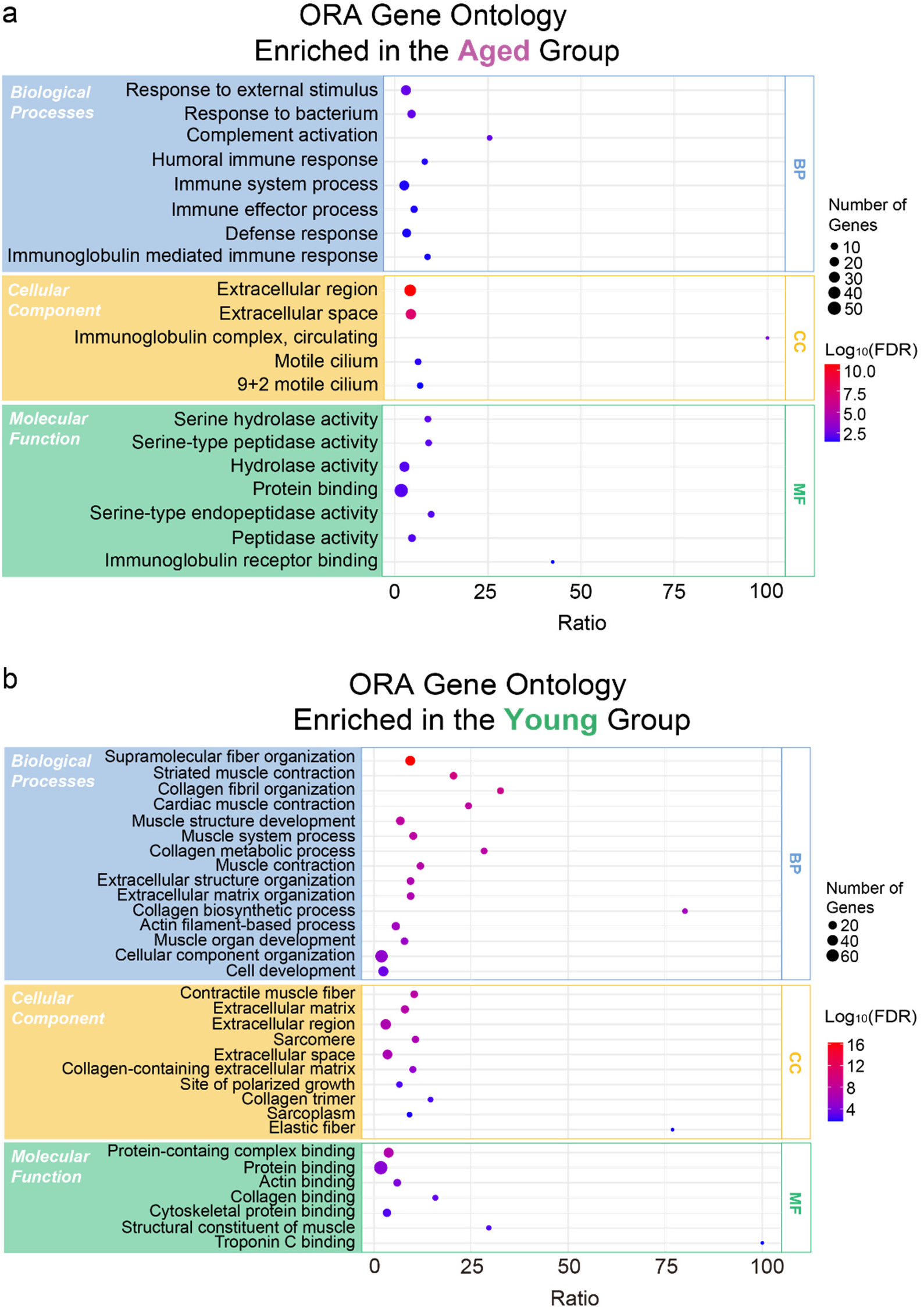
Over-representation analysis of differentially expressed proteins from single-factor aged versus young comparison. **a** Over-representation analysis (ORA) of proteins upregulated in the aged group. **b** ORA of proteins upregulated in the young group. Results are shown for Gene Ontology (GO) categories including Biological Process (BP), Cellular Component (CC), and Molecular Function (MF). The Reactome pathways were significantly enriched among differentially expressed proteins in the young group are shown in Fig. 2f. No Reactome pathways were significantly enriched among differentially expressed proteins in the aged group. Bubble size indicates the number of overlapping gene hits, and color scale represents –Log₁₀ (FDR).

**Fig. S4:**
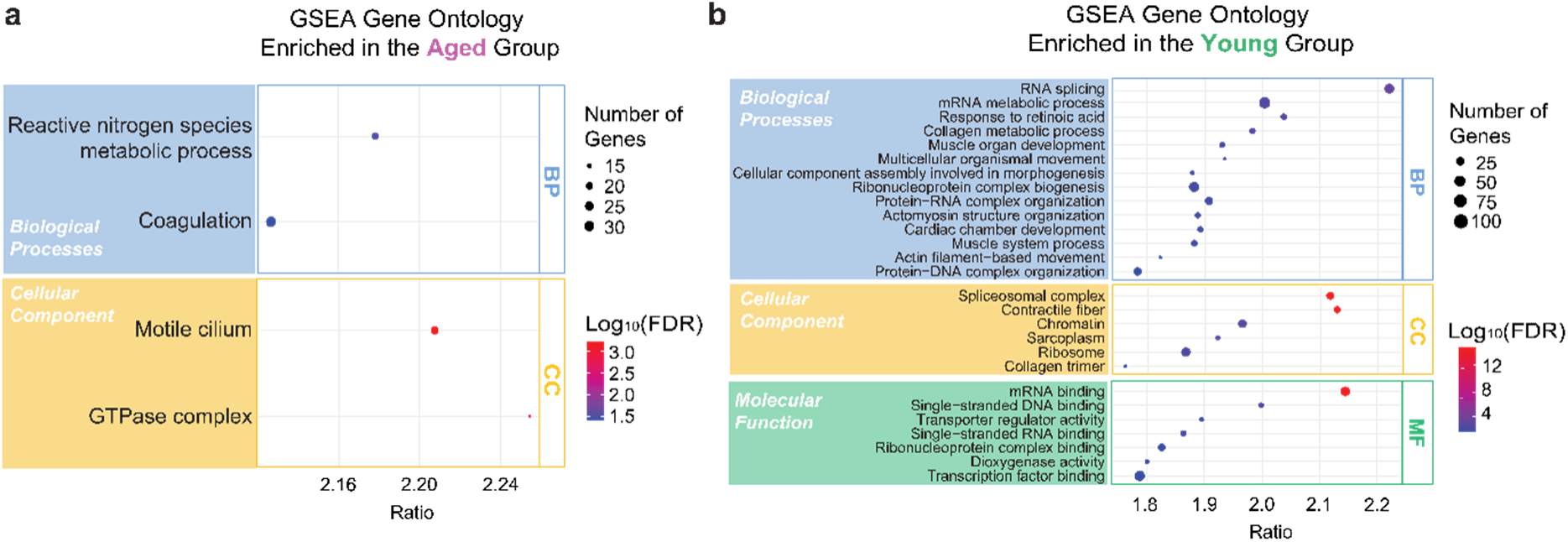
Gene set enrichment analysis of the full ranked proteome in single-factor aged versus young comparison. Gene set enrichment analysis (GSEA) of all quantified proteins ranked by Log₂(fold change) (aged versus young). **a** GSEA of proteins identified in the aged group. No significant terms were identified under Molecular Function. **b** GSEA of proteins identified in the young group. Bubble size indicates the number of leading-edge gene hits, and color represents –Log₁₀(FDR).

**Fig. S5:**
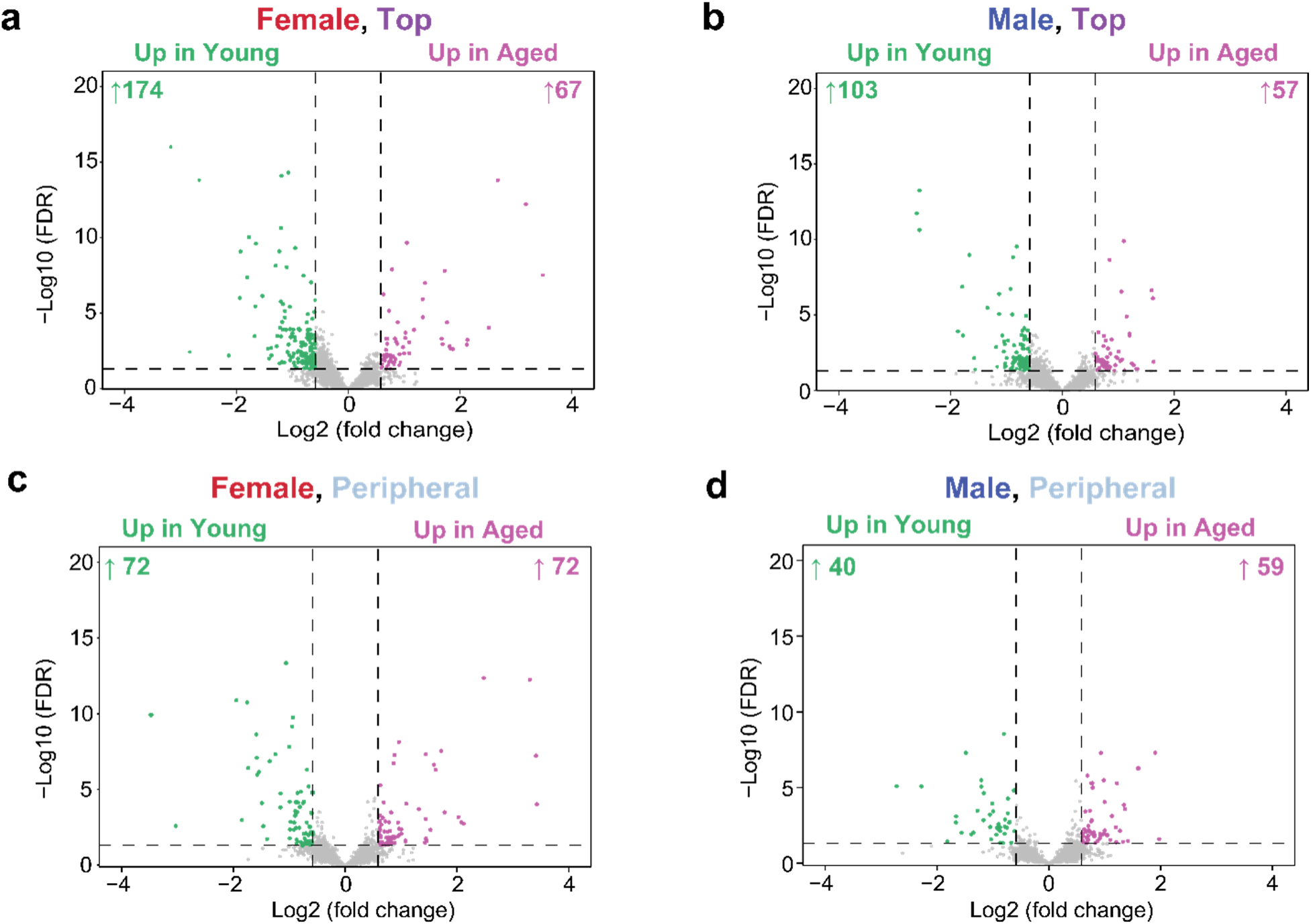
Multi-factor pairwise comparisons of aged versus young lungs in top and peripheral regions. Volcano plots show differential protein expression between aged and young rats for each sex and lung region: (**a**) female top, (**b**) male top, (**c**) female peripheral, and (**d**) male peripheral. Each point represents one quantified protein. Proteins with false discovery rate < 0.05 and |fold change| > 1.5 are considered significant. The number of upregulated proteins is indicated above each panel.

**Fig. S6:**
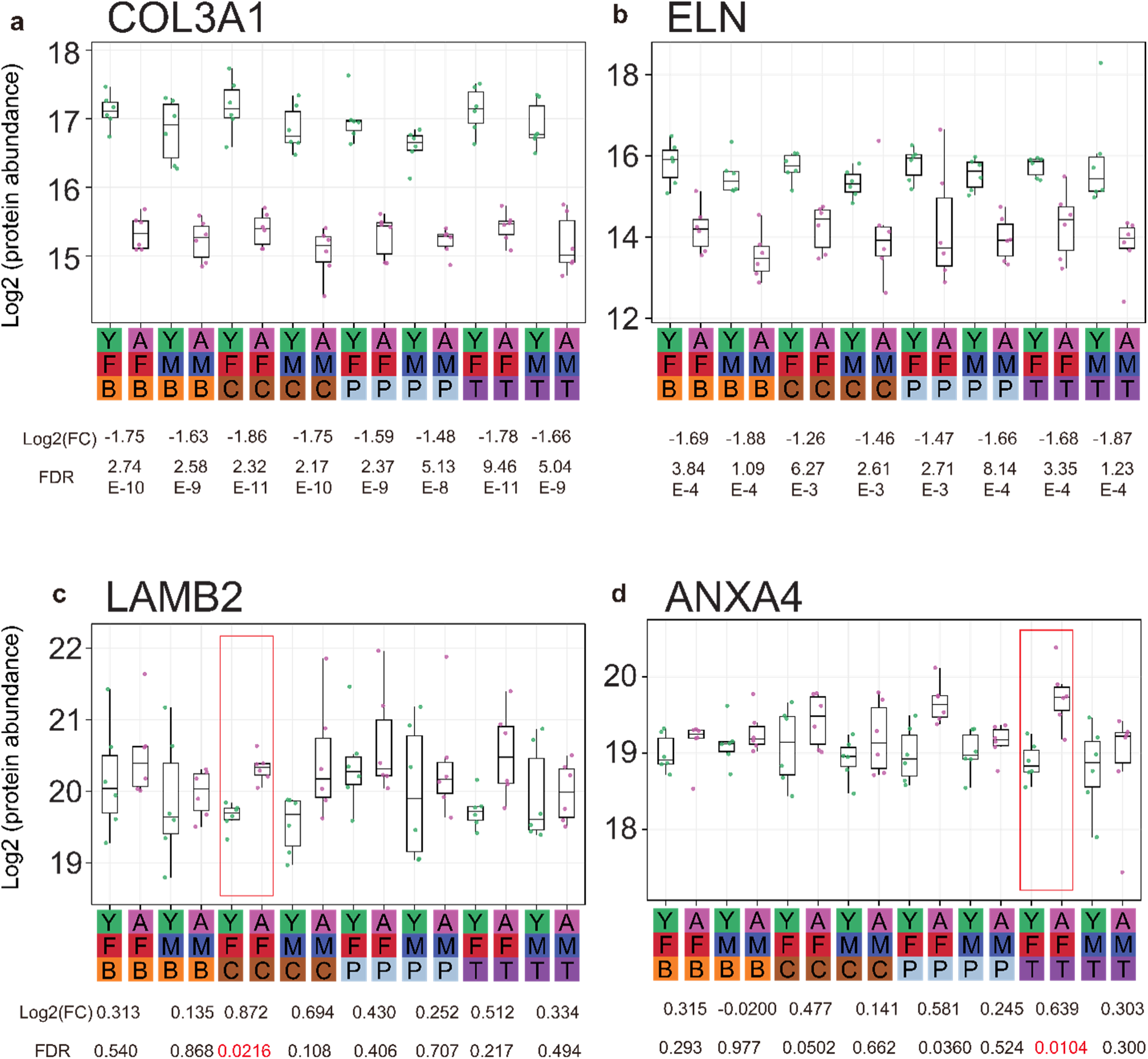
Representative boxplots of matrisome proteins exhibiting region and sex-specific differences in aged versus young comparison. **a** COL3A1 (Collagen alpha-1(III) chain), **b** ELN (Elastin), **c** LAMB2 (Laminin subunit beta-2), **d** ANAX4 (Annexin A4). Each point represents one biological replicate. There are n = 6 samples in each group. Log_2_(fold change) and false discovery rate (FDR) are listed below each aged versus young comparison. FDR < 0.05 comparisons are highlighted by a red rectangle.

**Fig. S7:**
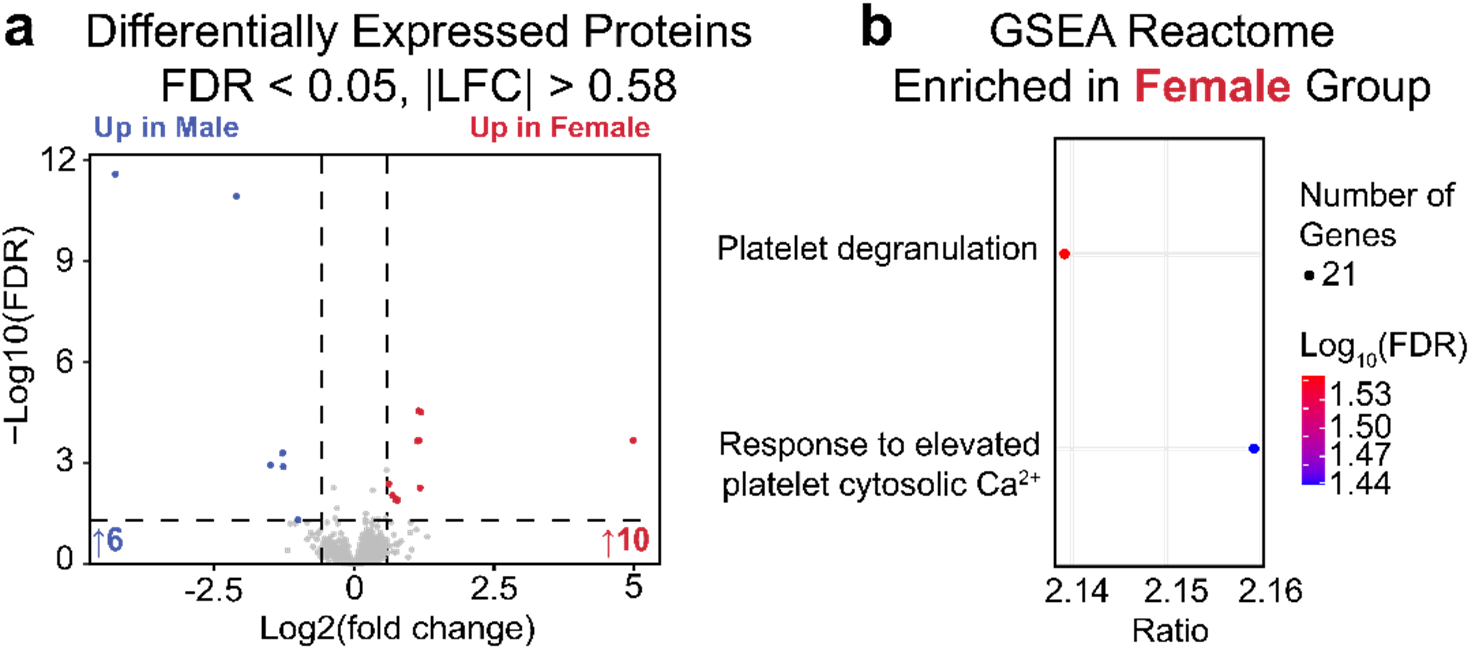
Differential proteomic analysis identifies female-enriched platelet degranulation and calcium signaling pathways. **a** Volcano plot of differentially expressed proteins between females and males across all samples. **b** Gene set enrichment analysis (GSEA) based on the ranked Log_2_(fold change) protein list (false discovery rate, FDR < 0.05). There are no female-enriched Gene Ontology (GO) categories in Biological Process (BP), Cellular Component (CC) or Molecular Function (MF). There are no male-enriched Reactome pathways or GO categories (FDR < 0.05).

**Fig. S8:**
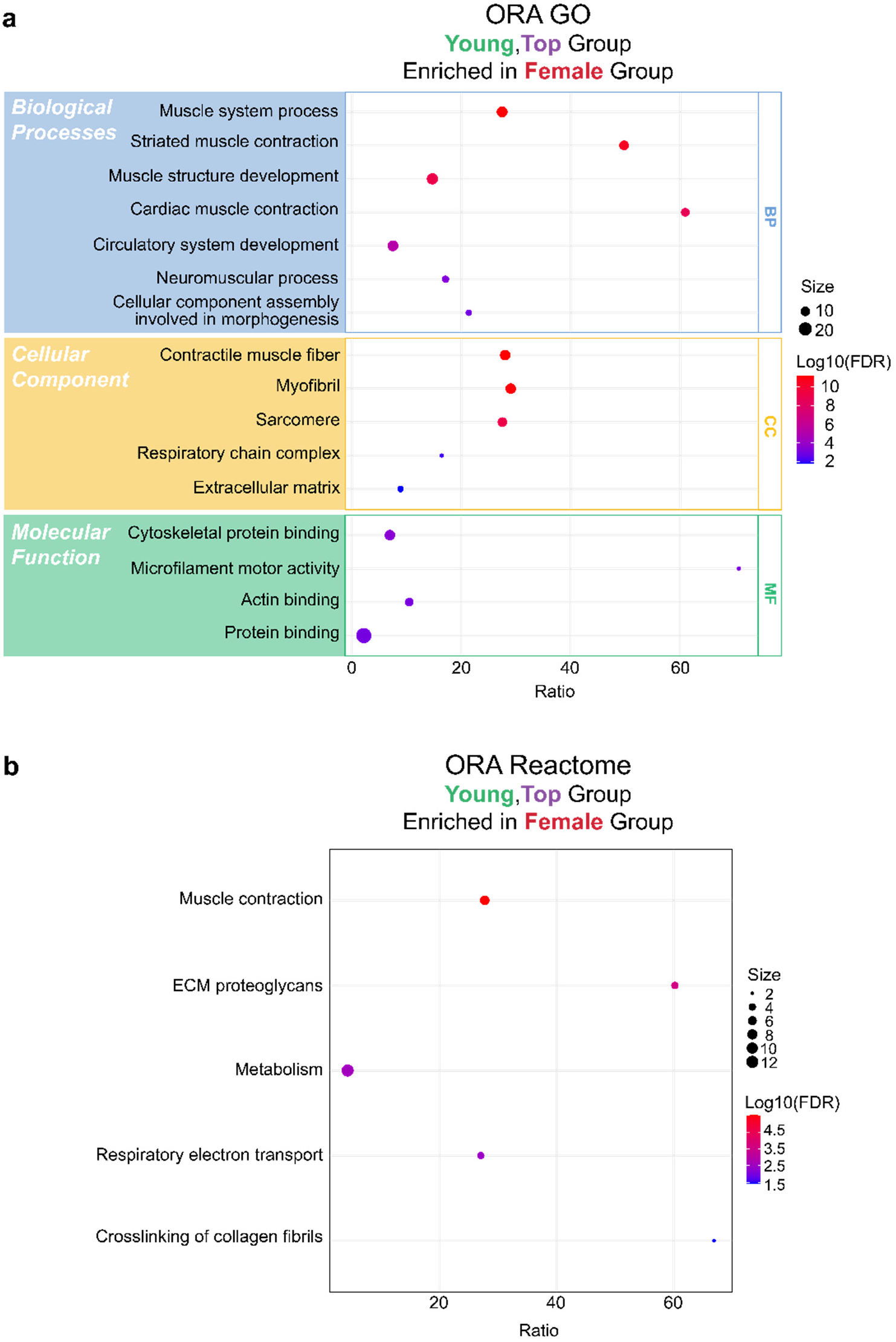
Over-representation analysis of differentially expressed proteins from female versus male comparison in top young region. **a** Gene Ontology (GO) enrichment by over-representation analysis (ORA) of proteins upregulated in the female group from female versus male comparison in top young region. **b** Reactome enrichment by ORA of proteins upregulated in the female group from female versus male comparison in top young region. Bubble size indicates the number of overlapping gene hits, and color scale represents –Log₁₀ (FDR).

**Fig. S9:**
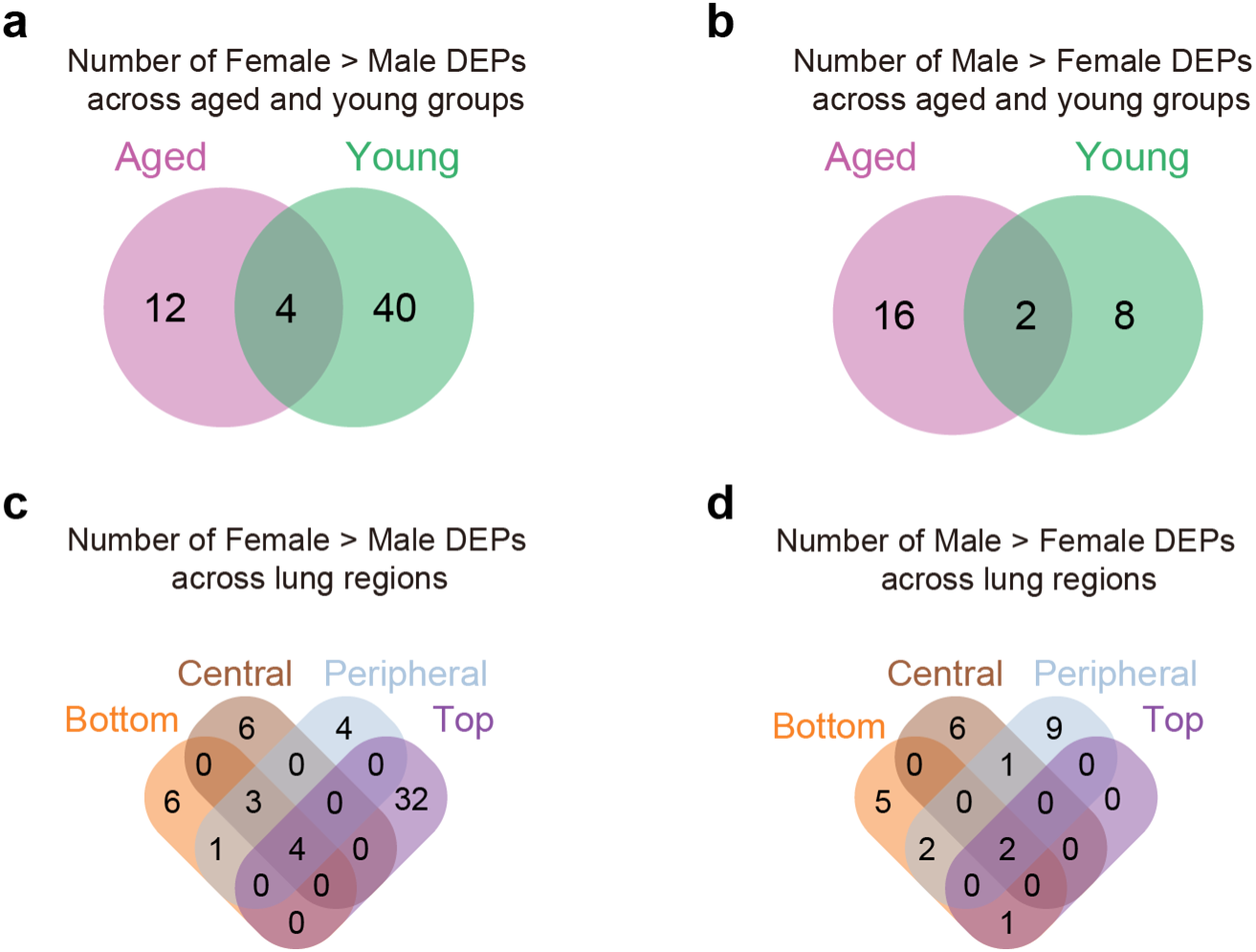
Venn diagrams of differentially expressed proteins from female versus male multi-factor comparisons in different age and region groups. **a** Venn diagram shows the overlap of differentially expressed proteins (DEPs) upregulated in female tissues in female versus male comparisons, grouped by age (aged, young). **b** Venn diagram shows the overlap of DEPs upregulated in male tissues in female versus male comparisons, grouped by age (aged and young). **c–d** Venn diagrams showing the overlap of DEPs between female and male samples across lung regions (bottom, central, peripheral, and top). Proteins are categorized into shared and region-specific groups. Detailed gene symbols corresponding to each subgroup are listed in Table S15.

**Fig. S10:**
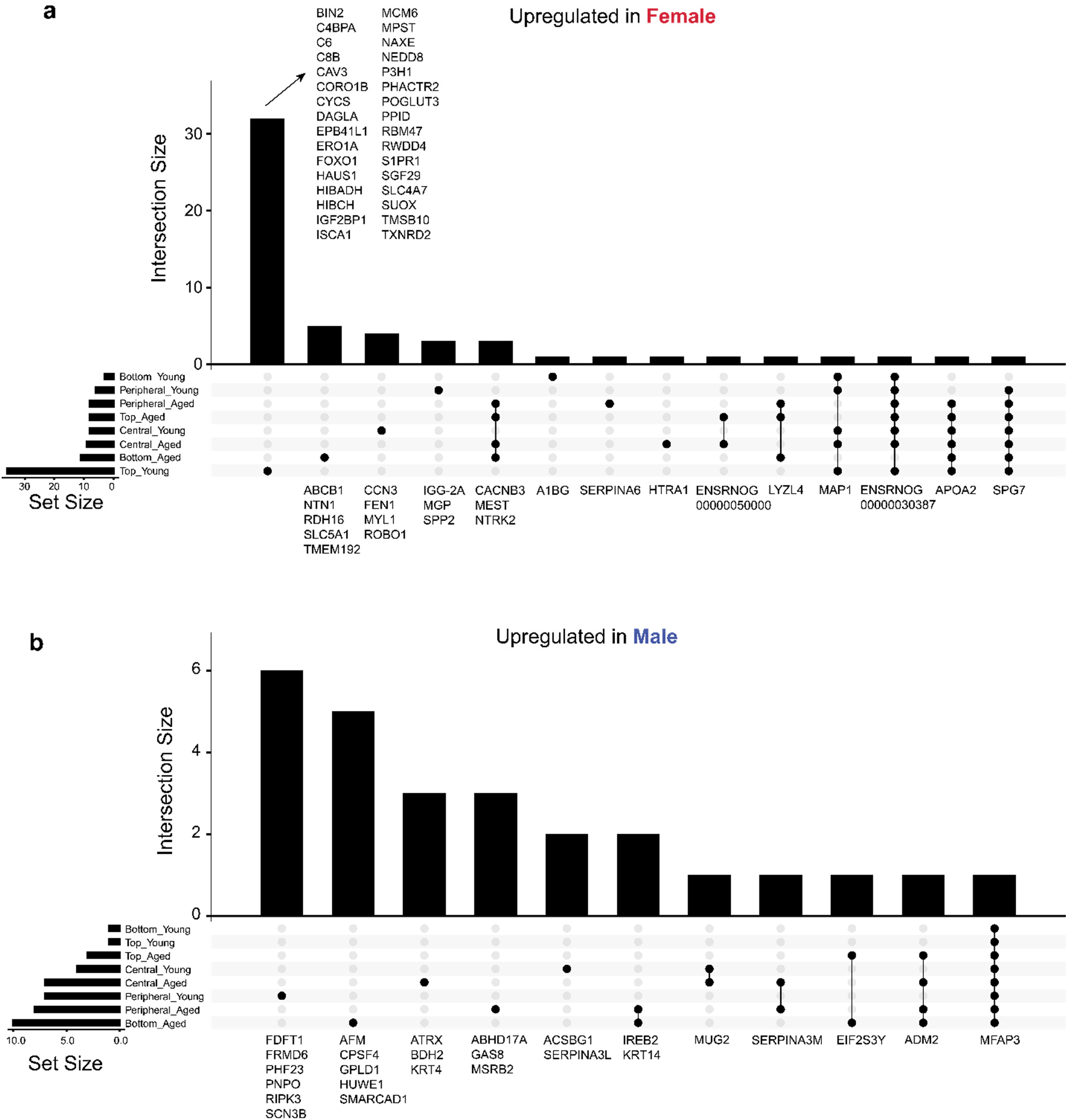
UpSet plots of differentially expressed proteins from female versus male multi-factor comparisons in different age and region groups. **a** UpSet plot shows the overlap of differentially expressed proteins (DEPs) upregulated in female tissues across eight pairwise female versus male comparisons, grouped by age (aged, young) and lung region (bottom, central, peripheral, top). **b** UpSet plot shows the overlap of DEPs upregulated in male tissues across the same comparisons. Gene symbols are labeled in each subgroup.

**Fig. S11:**
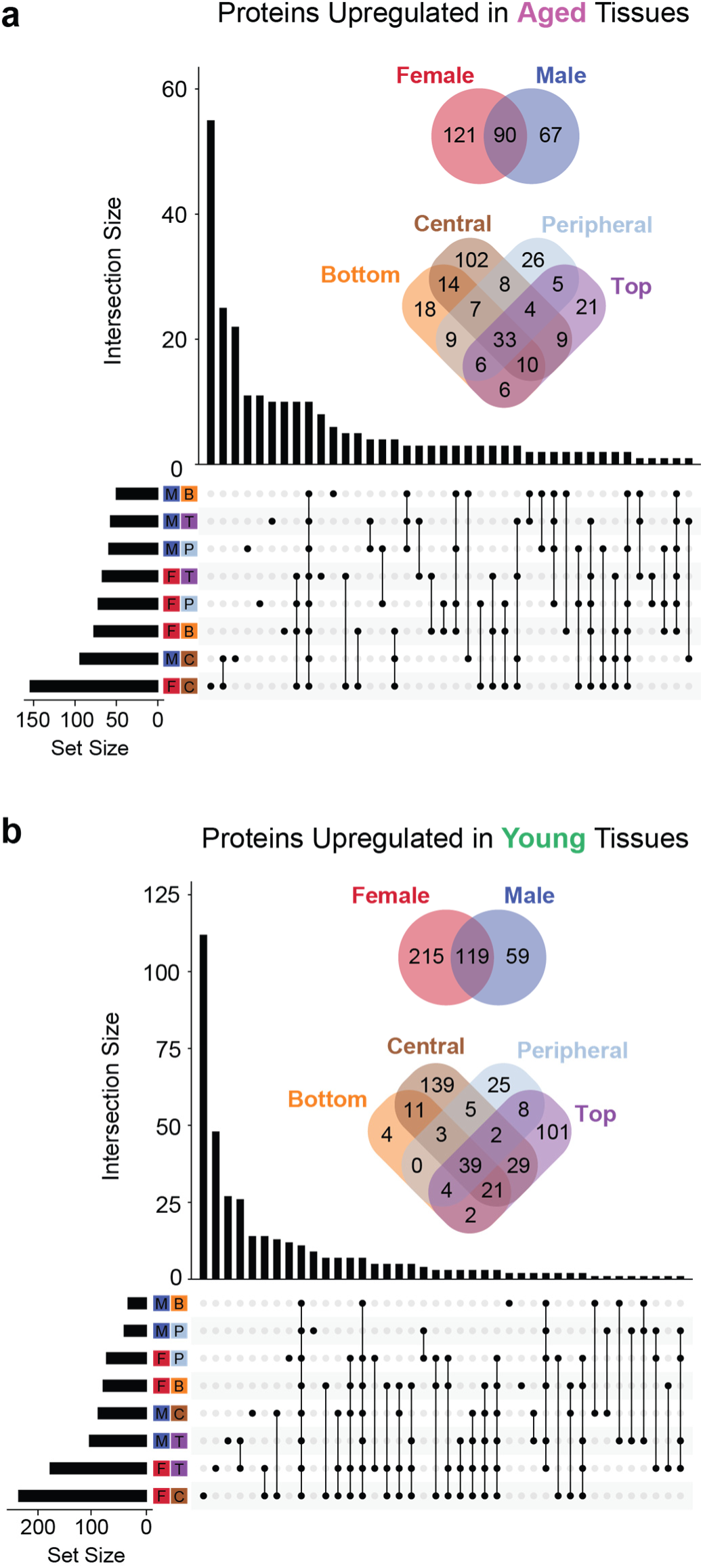
Overlap of differentially expressed proteins upregulated in aged and young tissues across multi-factor comparisons. **a** UpSet plot and Venn diagrams show the overlap of differentially expressed proteins (DEPs) upregulated in aged tissues across eight pairwise aged versus young comparisons, grouped by sex (female, male) and lung region (bottom, central, peripheral, top). **b** UpSet plot and Venn diagrams show the overlap of DEPs upregulated in young tissues across the same comparisons. Gene symbol lists are included in Supplementary Table 26.

**Fig. S12:**
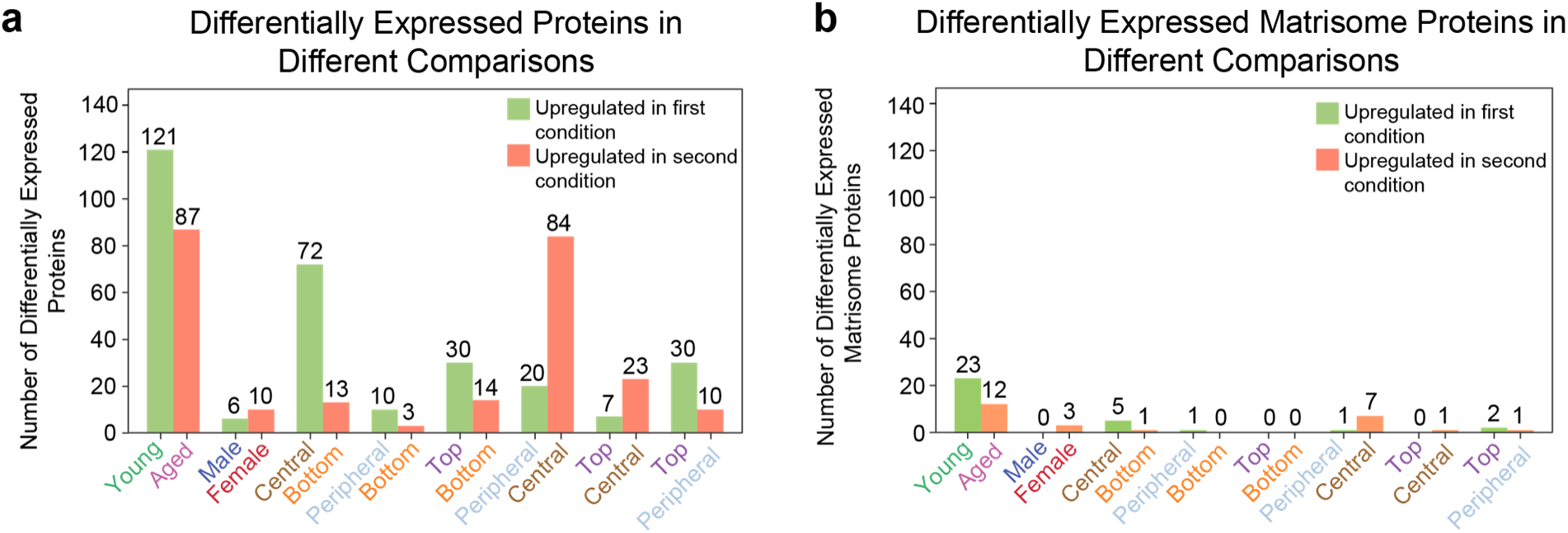
Summary of differentially expressed proteins from single-factor comparisons across age, sex, and lung regions. **a** Total number of differentially expressed proteins (DEPs) identified in each single-factor comparison, including age (young versus aged), sex (male versus female), and all regional pairs (bottom, central, peripheral, top). Bars indicate proteins upregulated in the first (green) or second (orange) condition, based on FDR < 0.05 and |fold change| > 1.5. **b** Corresponding summary for differentially expressed matrisome proteins.

**Fig. S13:**
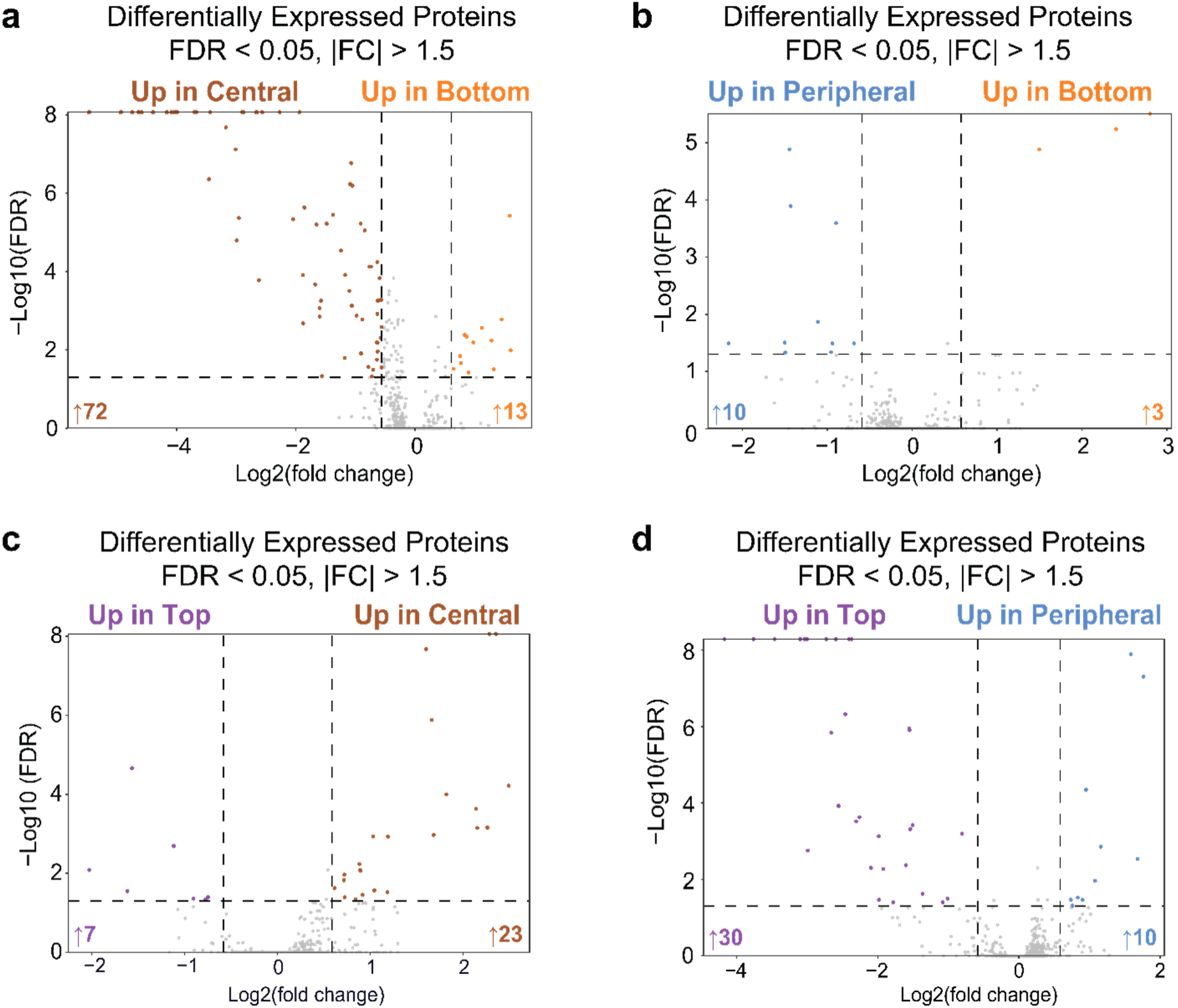
Single-factor regional comparisons of lung proteome. Volcano plots showing differential protein expression across lung regions not included in Fig. 6. Each point represents one quantified protein, with vertical lines marking |fold change| = 1.5 and horizontal lines indicating FDR = 0.05. (**a**) central versus bottom, (**b**) peripheral versus bottom, (**c**) top versus central, and (**d**) top versus peripheral comparisons. FDR, false discovery rate.

**Fig. S14:**
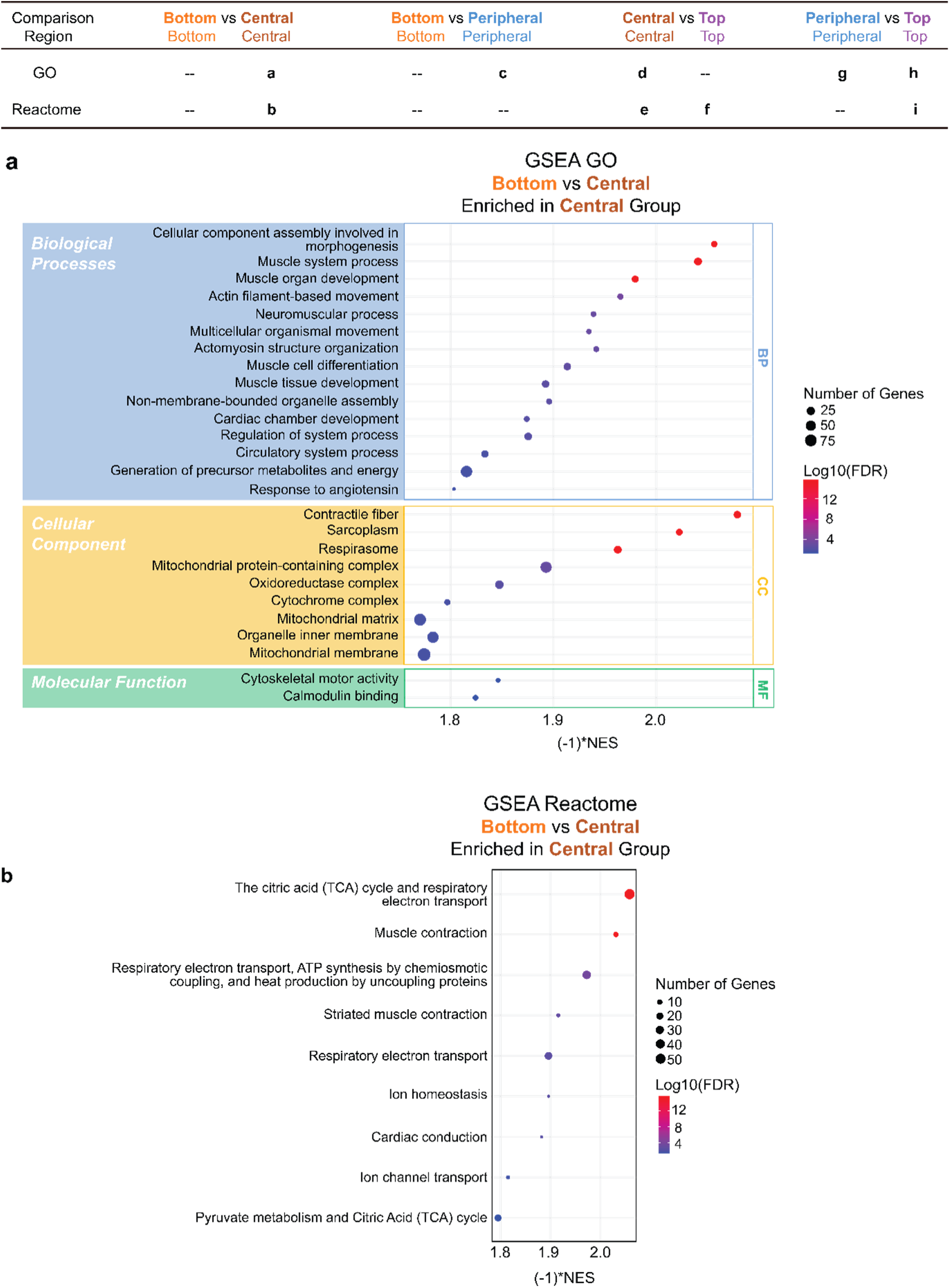
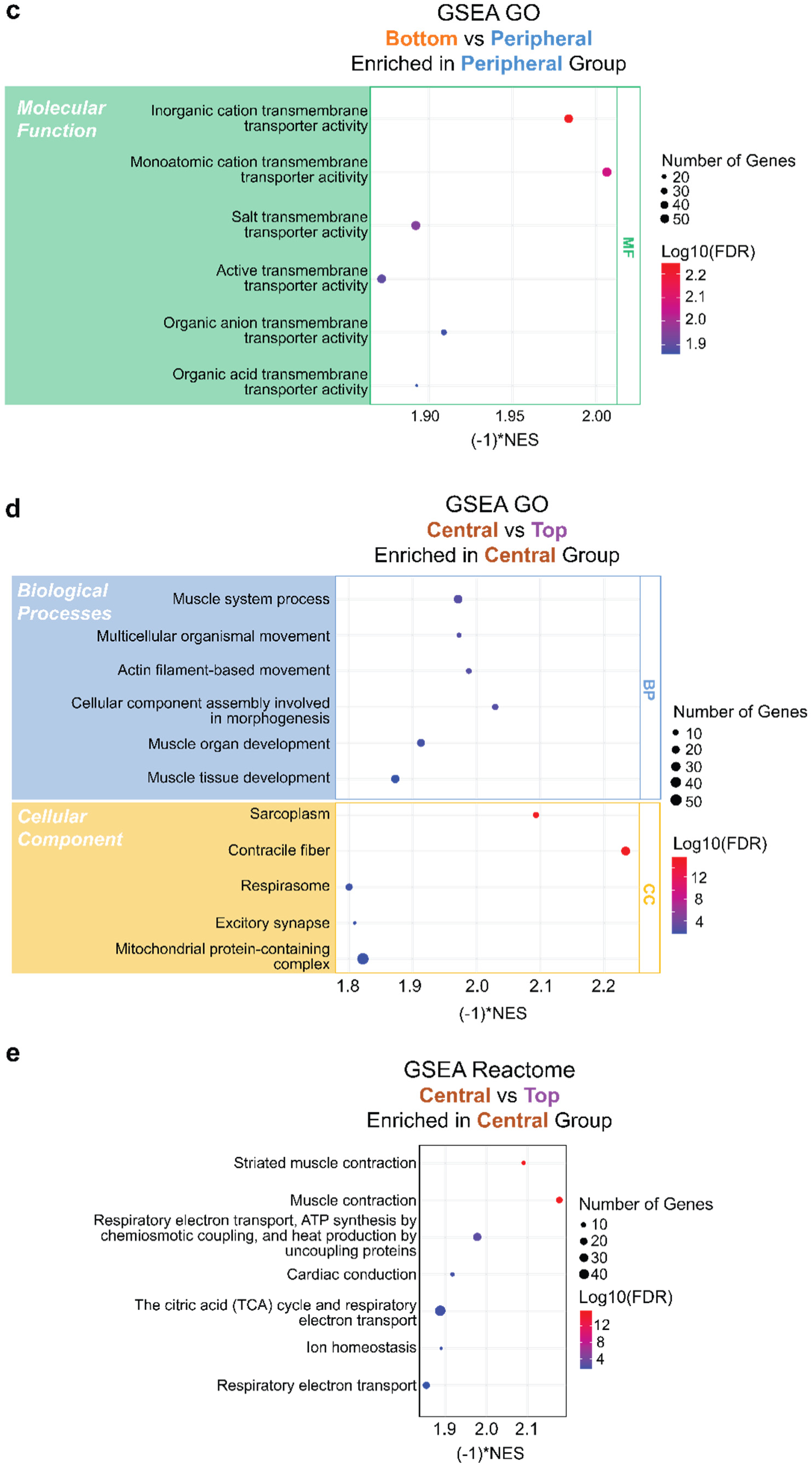
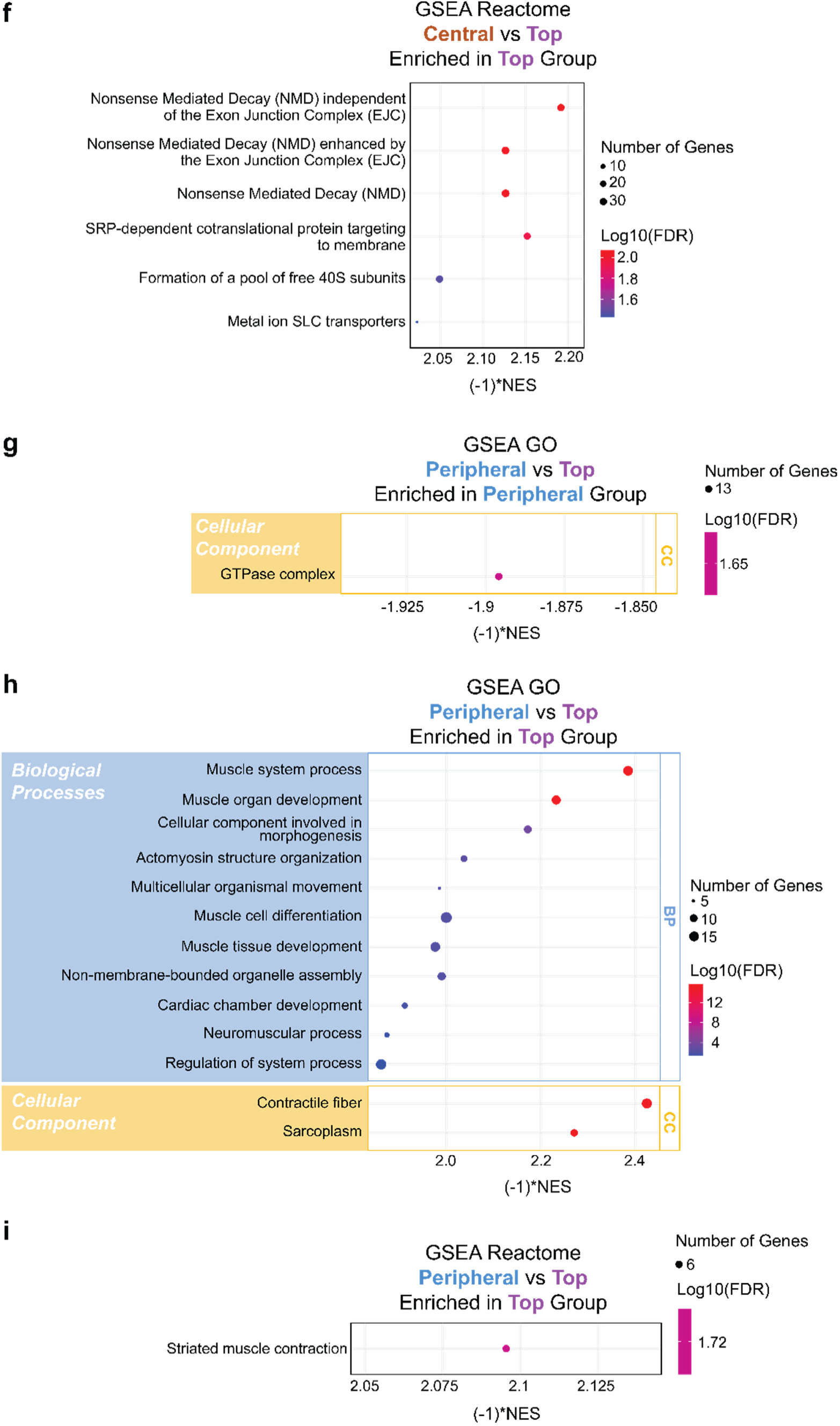
Gene set enrichment analysis of the full ranked proteome in the single-factor bottom versus central, bottom versus peripheral, central versus top, and peripheral versus top comparisons with significance. **a** Gene Ontology (GO) terms enriched in central region in bottom versus central comparison. **b** Reactome terms enriched in central region in bottom versus central comparison. **c** GO terms enriched in peripheral region in bottom versus peripheral comparison. **d** GO terms enriched in central region in central versus top comparison. **e** Reactome terms enriched in central region in central versus top comparison. **f** Reactome terms enriched in top region in central versus top comparison. **g** GO terms enriched in peripheral region in peripheral versus top comparison. **h** GO terms enriched in top region in peripheral versus top comparison. **i** Reactome terms enriched in top region in peripheral versus top comparison. Selected pathways with significance (FDR < 0.05) are displayed. Bubble size indicates the number of leading-edge gene hits, and color represents – Log₁₀(FDR). Results are shown for GO categories including Biological Process (BP), Cellular Component (CC), and Molecular Function (MF).

**Fig. S15:**
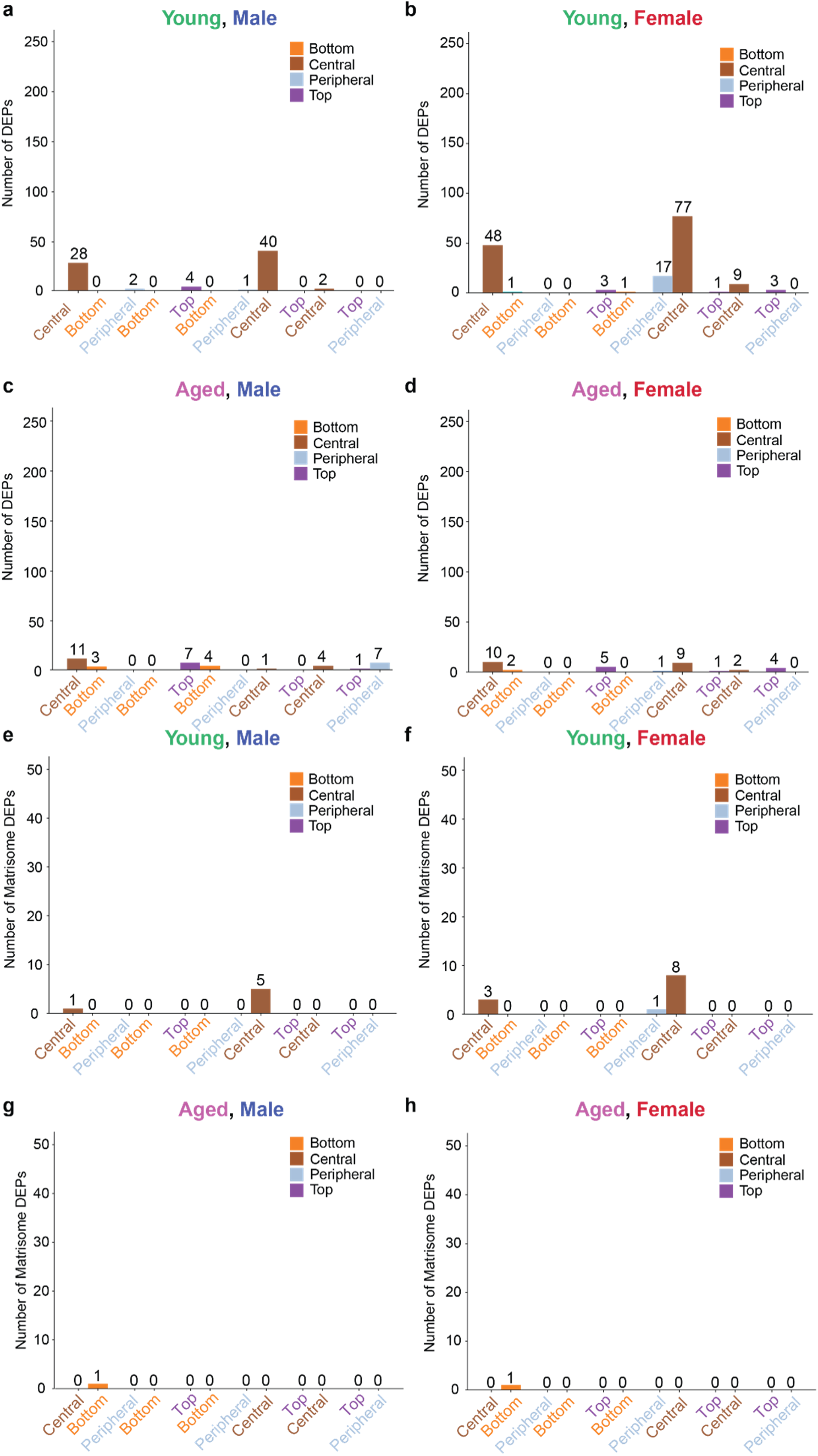
Multi-factor pairwise comparisons of regional proteomic differences across age and sex. **a–d** Number of differentially expressed proteins (DEPs) (|fold change| > 1, FDR < 0.05) identified in regional comparisons within each group: (a) young male, (b) young female, (c) aged male, and (d) aged female. Each two adjacent bar represents a regional pairwise comparison (e.g., central versus bottom, central versus peripheral). **e–h** Corresponding numbers of differentially expressed matrisome proteins for the same comparisons: (**e**) young male, (**f**) young female, (**g**) aged male, and (**h**) aged female.

**Fig. S16:**
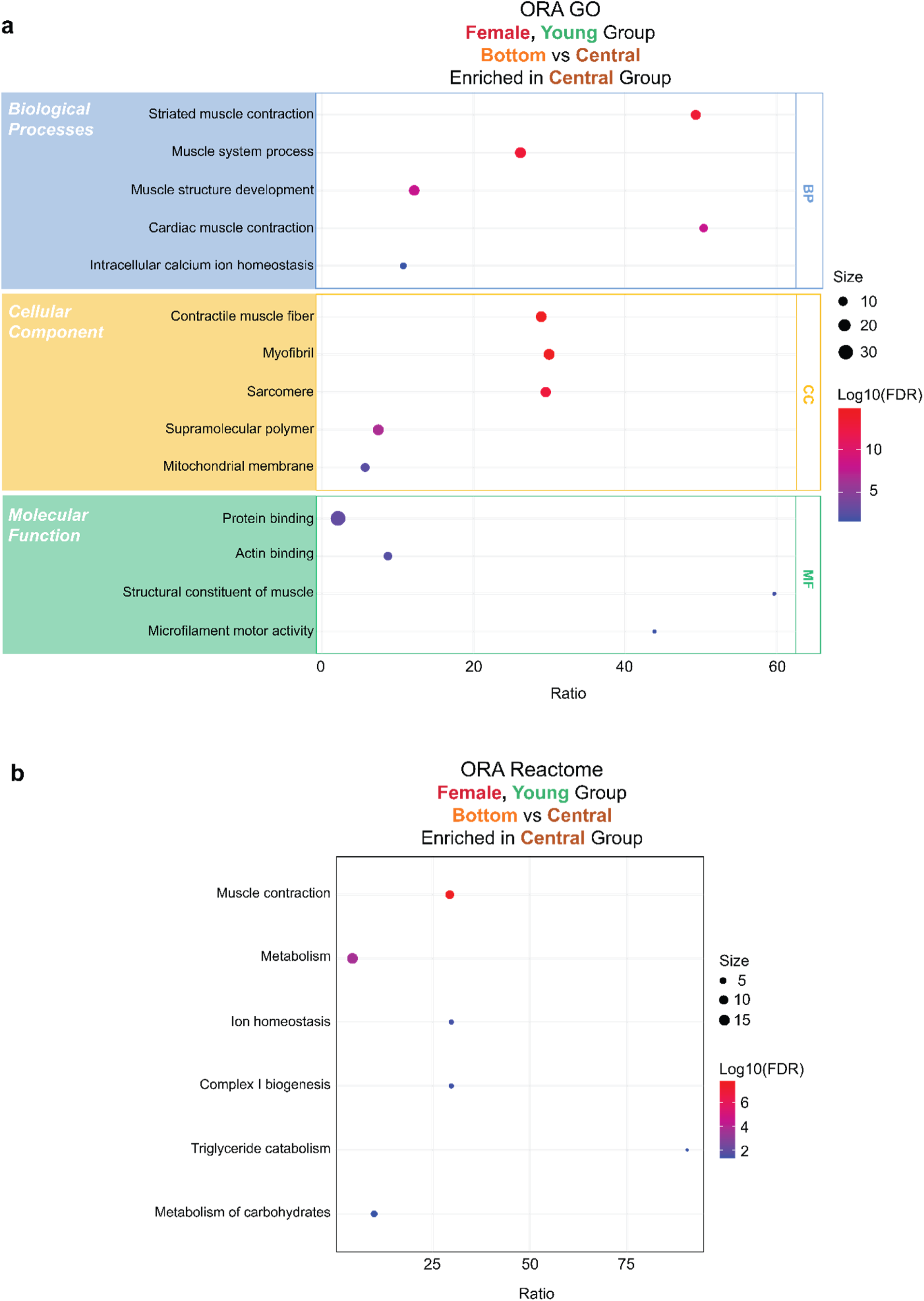
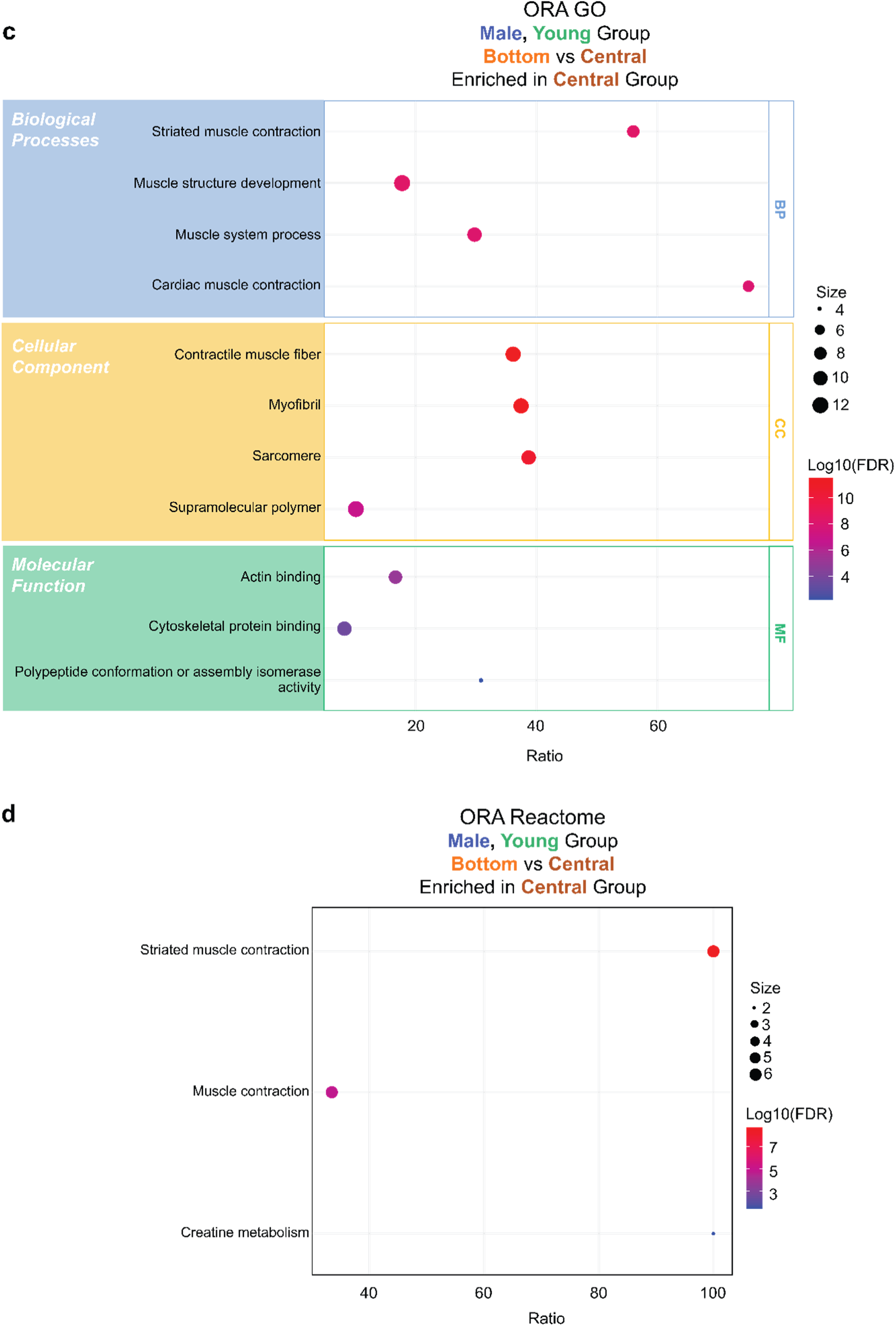
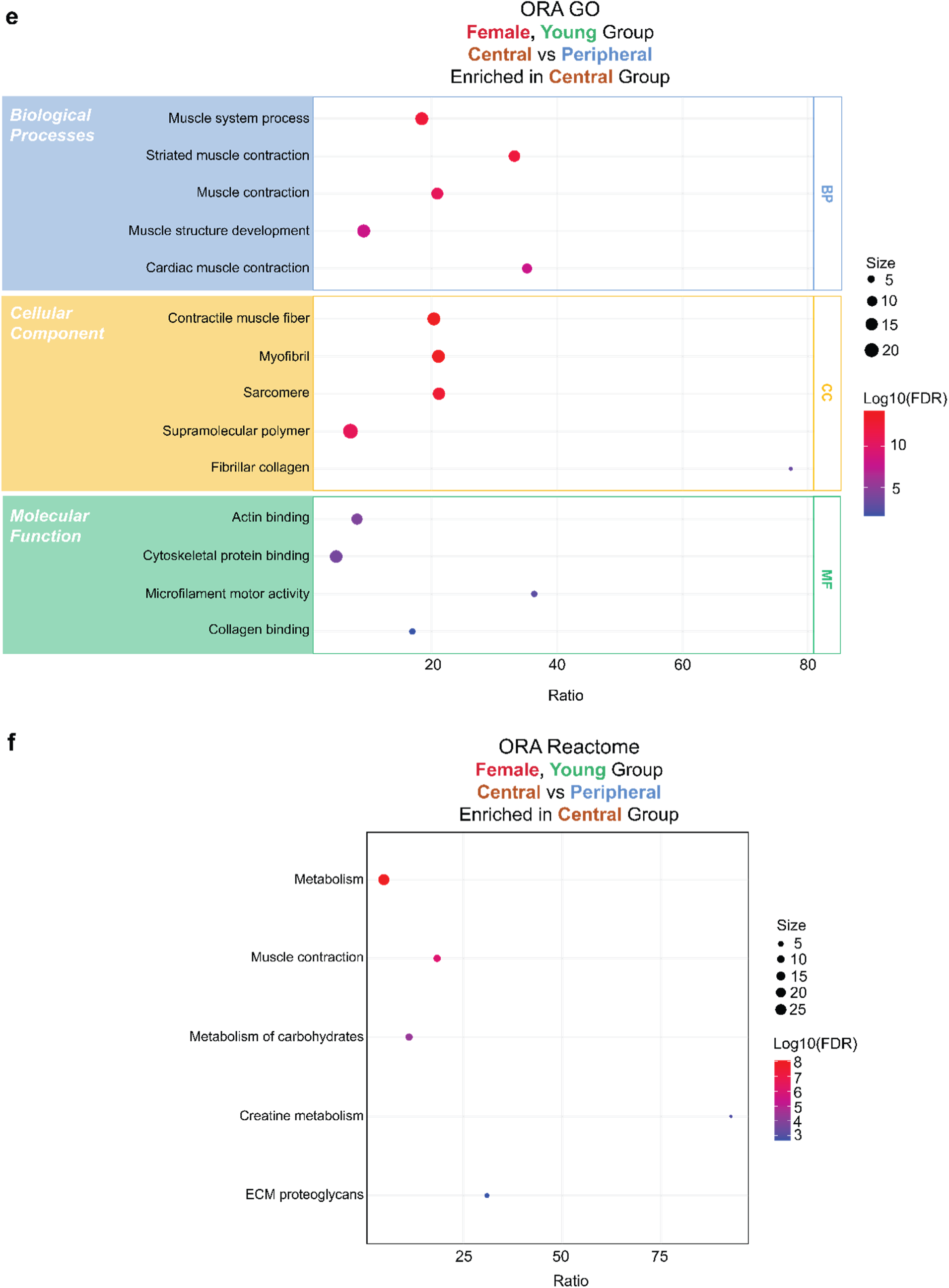
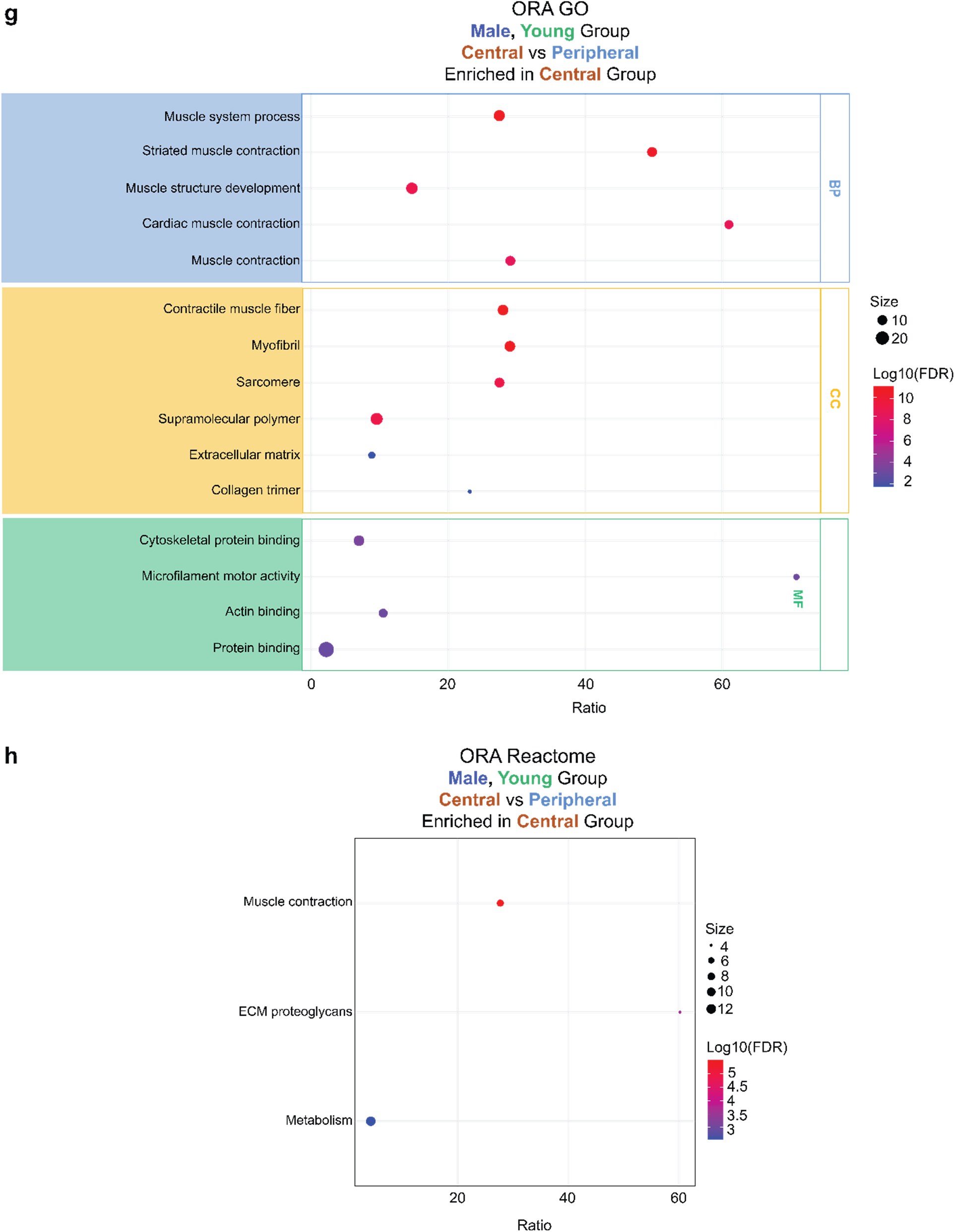
Over-representation analysis of differentially expressed proteins from central versus peripheral and central versus bottom in young female and young male groups. **a** Gene Ontology (GO) enrichment by over-representation analysis (ORA) of proteins upregulated in the central group from central versus bottom comparison in female young group. **b** Reactome enrichment by ORA of proteins upregulated in the central group from bottom versus central comparison in female young group. **c** GO enrichment by ORA of proteins upregulated in the central group from bottom versus central comparison in male young group. **d** Reactome enrichment by ORA of proteins upregulated in the central group from bottom versus central comparison in male young group. **e** GO enrichment by ORA of proteins upregulated in the central group from central versus peripheral comparison in female young group. **f** Reactome enrichment by ORA of proteins upregulated in the central group from central versus peripheral comparison in female young group. **g** GO enrichment by ORA of proteins upregulated in the central group from central versus peripheral comparison in male young group. **h** Reactome enrichment by ORA of proteins upregulated in the central group from peripheral versus bottom comparison in male young group. Selected pathways with significance (FDR < 0.05) are displayed. Bubble size indicates the number of overlapping gene hits, and color scale represents –Log₁₀ (FDR). Results are shown for GO categories including Biological Process (BP), Cellular Component (CC), and Molecular Function (MF).

**Fig. S17:**
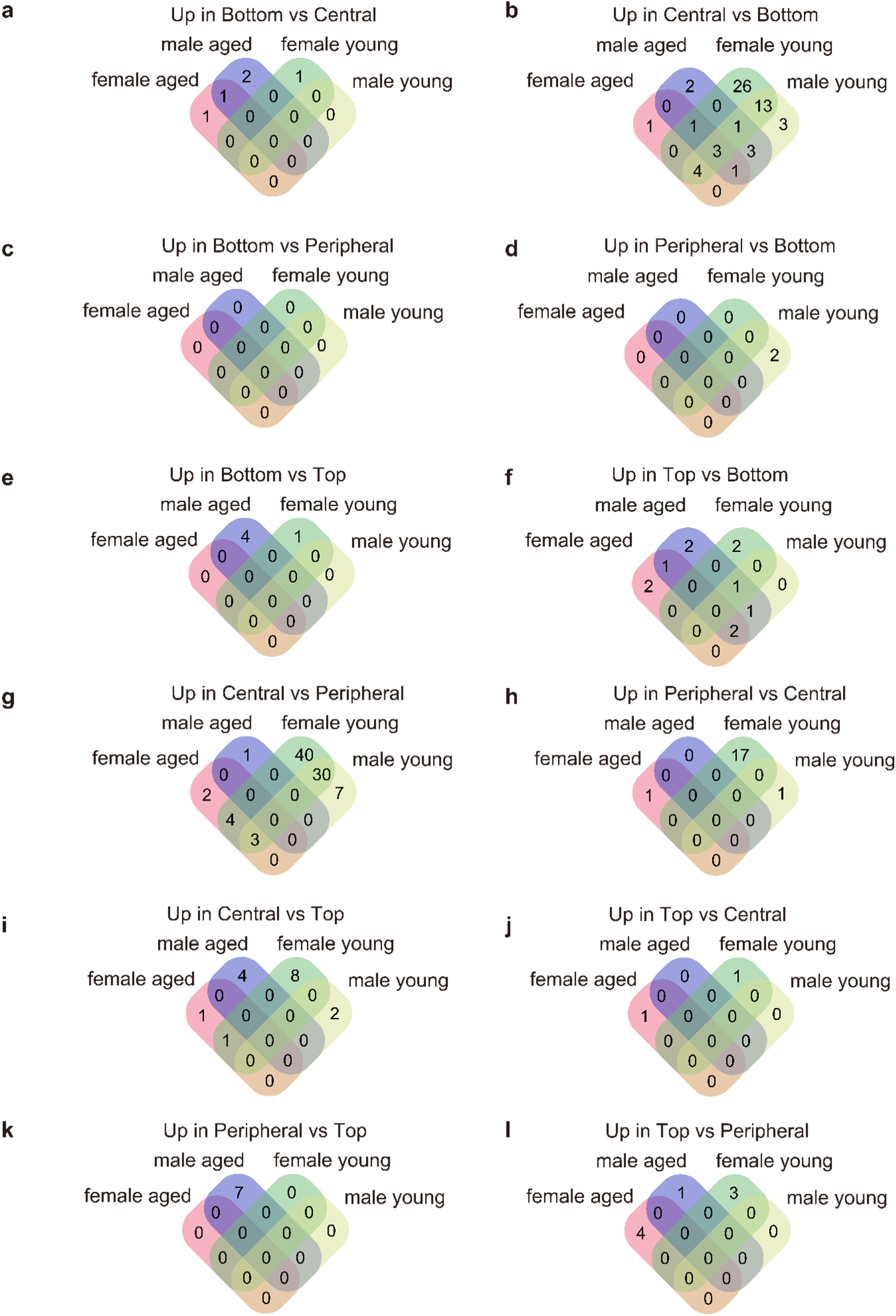
Venn diagrams of differentially expressed proteins from region versus region multi-factor comparisons in different age and sex groups. **a** Venn diagram shows the overlap of differentially expressed proteins (DEPs) upregulated in bottom tissues in bottom versus central comparisons, grouped by age (aged and young) and sex (female and male). **b** Venn diagram shows the overlap of DEPs upregulated in central tissues in bottom versus central comparisons, grouped by age (aged and young) and sex (female and male). **c** Venn diagram shows the overlap of DEPs upregulated in bottom tissues in bottom versus peripheral comparisons, grouped by age (aged and young) and sex (female and male). **d** Venn diagram shows the overlap of DEPs upregulated in peripheral tissues in bottom versus peripheral comparisons, grouped by age (aged and young) and sex (female and male). **e** Venn diagram shows the overlap of DEPs upregulated in bottom tissues in bottom versus top comparisons, grouped by age (aged and young) and sex (female and male). **f** Venn diagram shows the overlap of DEPs upregulated in top tissues in bottom versus top comparisons, grouped by age (aged and young) and sex (female and male). **g** Venn diagram shows the overlap of DEPs upregulated in central tissues in central versus peripheral comparisons, grouped by age (aged and young) and sex (female and male). **h** Venn diagram shows the overlap of DEPs upregulated in peripheral tissues in central versus peripheral comparisons, grouped by age (aged and young) and sex (female and male). **i** Venn diagram shows the overlap of DEPs upregulated in central tissues in central versus top comparisons, grouped by age (aged and young) and sex (female and male). **j** Venn diagram shows the overlap of DEPs upregulated in top tissues in central versus top comparisons, grouped by age (aged and young) and sex (female and male). **k** Venn diagram shows the overlap of DEPs upregulated in peripheral tissues in peripheral versus top comparisons, grouped by age (aged and young) and sex (female and male). **l** Venn diagram shows the overlap of DEPs upregulated in top tissues in peripheral versus top comparisons, grouped by age (aged and young) and sex (female and male). Detailed protein lists in each group of all panels are listed in Table S24.

**Fig: S18.**
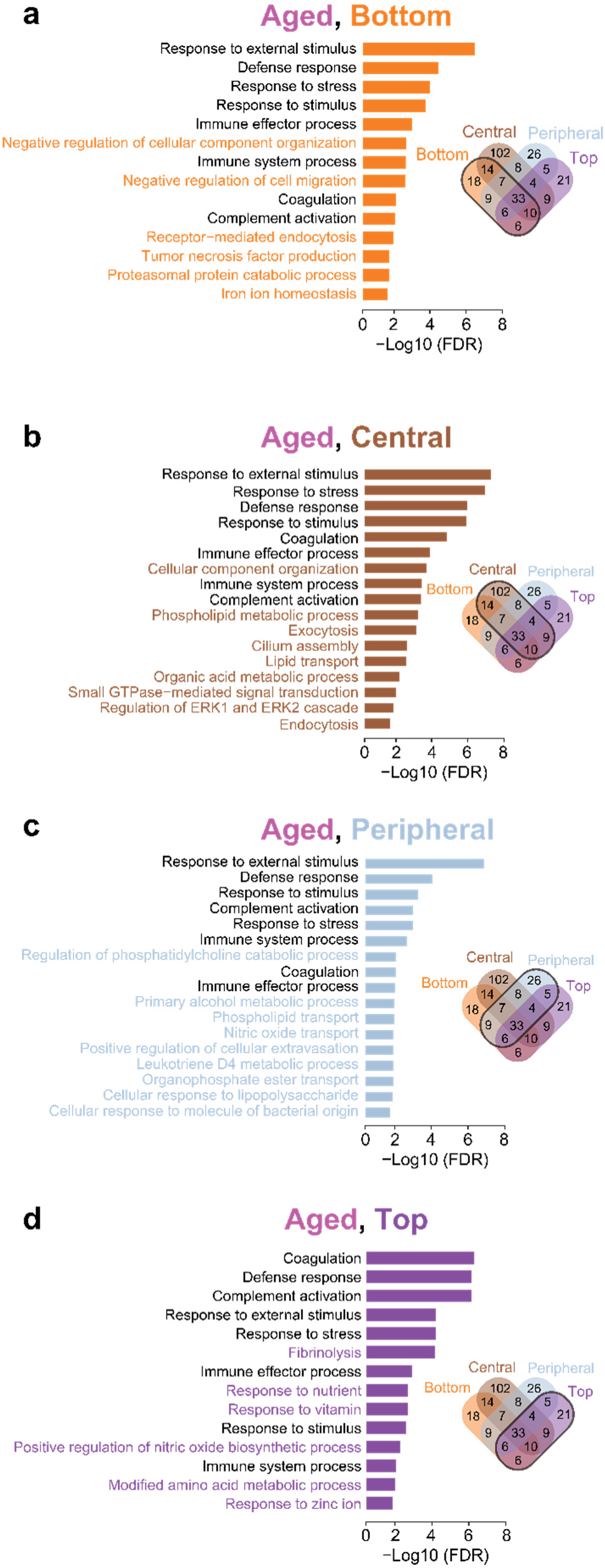
Region-specific over-representation analysis comparisons in aged tissues in aged versus young comparisons. Over-representation analysis (ORA) of proteins upregulated in aged (a–d) tissues across lung regions: (**a)** bottom; (**b**) central; (**c**) peripheral; (**d**) top. Upregulated proteins used for ORA input are highlighted in black circles from Venn diagrams (listed in Table S26). Bars represent significantly enriched gene ontology (GO) Biological Process (BP) terms, with pathway labels color-coded by region. Pathways shared with single-factor aged versus young comparisons are showsn in black, while region-specific pathways are highlighted in color. Selected pathways with significance (FDR < 0.05) are displayed.

**Fig: S19.**
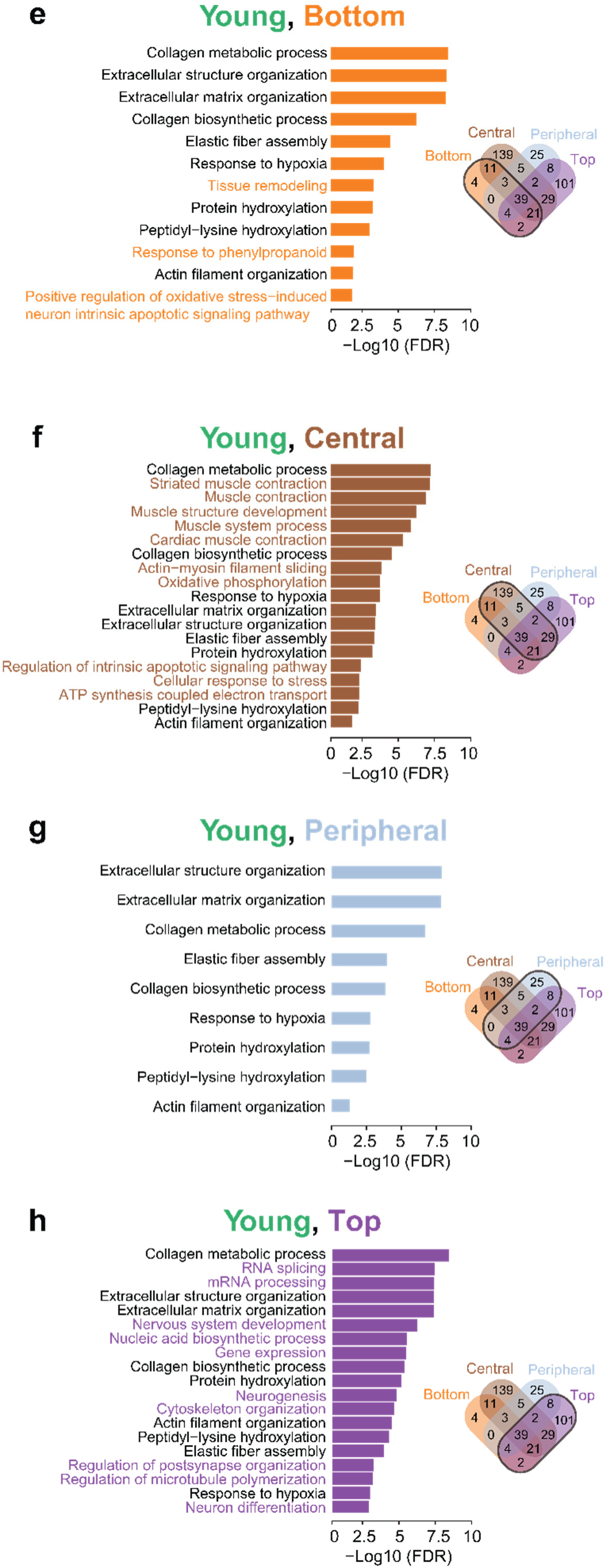
Region-specific over-representation analysis comparisons in young tissues in aged versus young comparisons. Over-representation analysis (ORA) of proteins upregulated in young (a–d) tissues across lung regions: (**a)** bottom; (**b**) central; (**c**) peripheral; (**d**) top. Upregulated proteins used for ORA input are highlighted in black circles from Venn diagrams (also listed in Table S26). Bars represent significantly enriched gene ontology (GO) Biological Process (BP) terms, with pathway labels color-coded by region. Pathways shared with single-factor aged versus young comparisons are shown in black, while region-specific pathways are highlighted in color. Selected pathways with significance (FDR < 0.05) are displayed.

